# Norepinephrine enhances oligodendrocyte precursor cell calcium dynamics in the cerebral cortex during arousal

**DOI:** 10.1101/2022.08.25.505119

**Authors:** Tsai-Yi Lu, Priyanka Hanumaihgari, Eric T. Hsu, Amit Agarwal, Dwight E. Bergles

## Abstract

Oligodendrocytes are generated from a widely distributed population of progenitors that express neurotransmitter receptors, but the mechanisms that alter activity of these oligodendrocyte precursor cells (OPCs) *in vivo* have not been determined. We generated a novel line of transgenic mice to express membrane-anchored GCaMP6s in OPCs and used longitudinal two-photon microscopy to monitor their Ca^2+^ changes in the cerebral cortex of awake mice. OPCs exhibited high rates of spontaneous activity, consisting of focal, transient Ca^2+^ increases within their highly ramified processes. Unexpectedly, these events occurred independent of excitatory neuron activity, but were inhibited by anesthesia, sedative agents, and antagonists of noradrenergic signaling. These norepinephrine enhanced Ca^2+^ dynamics rapidly declined as with differentiation. Selective knockout of α_1A_ adrenergic receptors in OPCs suppressed both spontaneous and locomotion-induced Ca^2+^ increases, indicating that OPCs are directly modulated by norepinephrine *in vivo*, providing a means to alter their dynamics and lineage progression during distinct brain states.

## Introduction

The adult central nervous system (CNS) contains an abundant population of glial progenitor cells that retain the ability to differentiate into oligodendrocytes. The continued presence of these lineage restricted glial cells^1^, which maintain a highly ordered, grid-like distribution in both white and gray matter, provides a means to generate new oligodendrocytes in response to changes in life experience^2–5^. These progenitors can also be mobilized to regenerate oligodendrocytes lost through injury or diseases such as multiple sclerosis^2^ and amyotrophic lateral sclerosis^1^. Although these glial cells have been justly termed oligodendrocyte precursor cells (OPCs), they are found in brain regions devoid of myelin^3^, they continually survey their environment using dynamic filopodia that extend from their fine, highly ramified processes^4^,and they migrate to sites of tissue injury like microglia to contribute to glial scar/barrier formation^4,5^. Moreover, recent studies indicate that OPCs can engulf cellular debris^6^, present exogenous antigen through MHC II^7,8^, and prune axons and synapses during development^6^, suggesting that they may have broader roles in tissue surveillance and homeostasis. In support of this hypothesis, genetic depletion of these cells from the CNS results in abnormal synaptic structure^9,10^, resulting in impaired sensory information processing^10^ and homeostatic control of metabolism^9^. However, the mechanisms used to control the behavior of these ubiquitous progenitors within neural circuits remain poorly understood.

Physiological studies of OPCs in culture and in *ex vivo* tissue slices indicate that they express a diverse array of neurotransmitter receptors and Ca^2+^ permeable ion channels^11–13^, suggesting that they are subject to acute neuromodulation. Transcriptional mRNA profiling at the bulk and single cell level support this hypothesis and have further expanded the range of potential receptors expressed by these ubiquitous progenitors^14–17^. Indeed, OPCs throughout the CNS form direct synapses with neurons^18,19^, in which they serve as a postsynaptic target, enabling transient activation of ionotropic glutamate or GABA_A_ receptors in their processes^18–20^. Activation of these receptors can influence their proliferation, differentiation, and response to injury, providing means to link neural activity to proliferation and lineage progression^21,22^. In addition, *in vitro* screens in both mouse and human OPCs have shown that their dynamics can be influenced by compounds that act on ligand gated receptors, including neuropeptide and acetylcholine receptors^23,24^. Manipulation of these receptors in OPCs *in vivo* can exert similar changes in lineage progression^23–25^, indicating that they can respond to neuromodulators in the intact CNS. However, despite this extensive phenotypic analysis, we know little about the activity patterns exhibited by OPCs in the intact brain under physiological conditions, the mechanisms that control signaling with these cells, or the behavioral states in which these cells become activated.

In contrast to most other cells in the adult CNS, OPCs can exist in many different states as they progress through the stages of cell division, transform into premyelinating oligodendrocytes, or activate apoptotic pathways^4^, highlighting the need to relate activity patterns to cell behavior. Recent *in vivo* imaging studies in the developing spinal cord of larval zebrafish revealed that OPCs exhibit periodic increases in intracellular Ca^2+^ that varied depending on soma location^26^, with cell activity higher in sparsely myelinated regions^26^. Moreover, *in vivo* imaging in anesthetized mice showed that activation of olfactory neurons with odorants results in Ca^2+^ increases within OPCs in activated glomeruli within the olfactory bulb^27^, a phenomenon that also appears to be independent of myelination. These results raise the possibility that OPCs can exist in distinct physiological states and exhibit regional differences in their response to neuronal activity. To define the mechanisms used to control the activity of OPCs *in vivo* during different behavioral states and at distinct cell stages, we generated a novel transgenic mouse line to enable visualization of dynamic Ca^2+^ signals within OPC processes and performed, time-lapse, two-photon (2P) imaging in the cerebral cortex of awake mice. Longitudinal imaging of OPCs revealed that they exhibit frequent focal Ca^2+^ increases within their processes that rapidly declined with lineage progression. Although OPCs receive synaptic input, this spontaneous activity was unexpectedly independent of local neuronal activity and ionotropic receptor activation. Instead, this local Ca^2+^ signaling was enhanced during periods of arousal, reflecting norepinephrine dependent activation of α_1_ adrenergic receptors in OPCs. These results indicate that OPCs are a direct target of norepinephrine, which provides a means to alter the physiology of these ubiquitous progenitors during distinct brain states.

## Results

### Generation of conditional mGCaMP6s mice to enable *in vivo* detection of OPC Ca^2+^ signaling

OPCs extend fine radial processes that ramify extensively within the surrounding neuropil and extend dynamic filopodia, providing a means to sense the release of neurotransmitters and neuromodulators^18,20,28^. However, the small amount of cytoplasm in these compartments make it difficult to achieve sufficient sensor concentration to resolve signaling events linked to changes in intracellular Ca^2+^. To examine Ca^2+^ changes within these small compartments *in vivo*, we generated a novel transgenic mouse line that enables conditional expression of a membrane anchored form of the genetically encoded calcium indicator GCaMP6s^29^ (*Rosa26-lsl-mGCaMP6s* mice, *mGCaMP6s*) (Extended Data Fig. 1a), which were then bred to *PDGFRα-CreER* BAC transgenic mice^1^ to express mGCaMP6s in OPCs. Administration of tamoxifen (TAM) to young adult *PDGFRα-CreER;mGCaMP6s* mice resulted in mGCaMP6s expression in the majority of OPCs, as assessed by co-localization between NG2 and GFP in brain sections prepared from these mice (Extended Data Fig. 1b-e). Comparison of mGCaMP6s expression in these mice with cytosolic GCaMP6s in *PDGFRα-CreER;R26-lsl-GCaMP6s* mice^29,30^, revealed that mGCaMP6s was more abundant than cytosolic GCaMP6s in the terminal branches of their processes (Extended Data Fig. 1f,g), suggesting that it may improve resolution of signaling events within these narrow compartments.

### OPCs exhibit transient Ca^2+^ increases in their processes *in vivo*

To assess whether OPCs in the mammalian brain exhibit dynamic Ca^2+^ signaling, we implanted chronic cranial windows over the visual cortex of young adult *PDGFRα-CreER;mGCaMP6s* mice (see Methods) and performed 2P fluorescence imaging under awake conditions, several weeks after administering TAM and habituating the mice to a head restrained imaging platform (Fig. 1a). Time lapse 2P imaging of individual OPCs in layer I of area V1 revealed that OPCs experienced frequent local increases in mGCaMP6s fluorescence in their processes, indicative of local elevation of intracellular Ca^2+^ (Fig. 1b, c, Supplementary Video 1, 2). These Ca^2+^ transients were highly variable in amplitude, time course, location, and size, and often propagated within an individual process (Fig. 1d). To quantify this dynamic behavior, we adapted <u>A</u>strocyte <u>Qu</u>antitative <u>A</u>nalysis (AQuA) software^31^ (see Methods) to detect and define the properties of each Ca^2+^ transient. In animals that were quietly resting on the stage, this activity was remarkably consistent, without obvious changes in frequency or duration, visible in raster plots of events ordered by time of occurrence (Fig. 1e). This activity was distributed throughout the cell, with Ca^2+^ transients observed in all visible processes within the imaging plane (Fig. 1f). Ca^2+^ transients occurred with an average frequency of 37 events/min within the imaging volume, with each event lasting for ~ 6 s (Extended Data Fig. 2a). In comparison to activity recorded in mice that expressed cytosolic GCaMP6s, mGCaMP6s-detected events were more frequent and larger in area (Extended Data Fig. 2a), supporting the hypothesis that membrane tethering GCaMP enhances the ability to resolve Ca^2+^ changes within the fine processes of OPCs.

**Figure 1.**
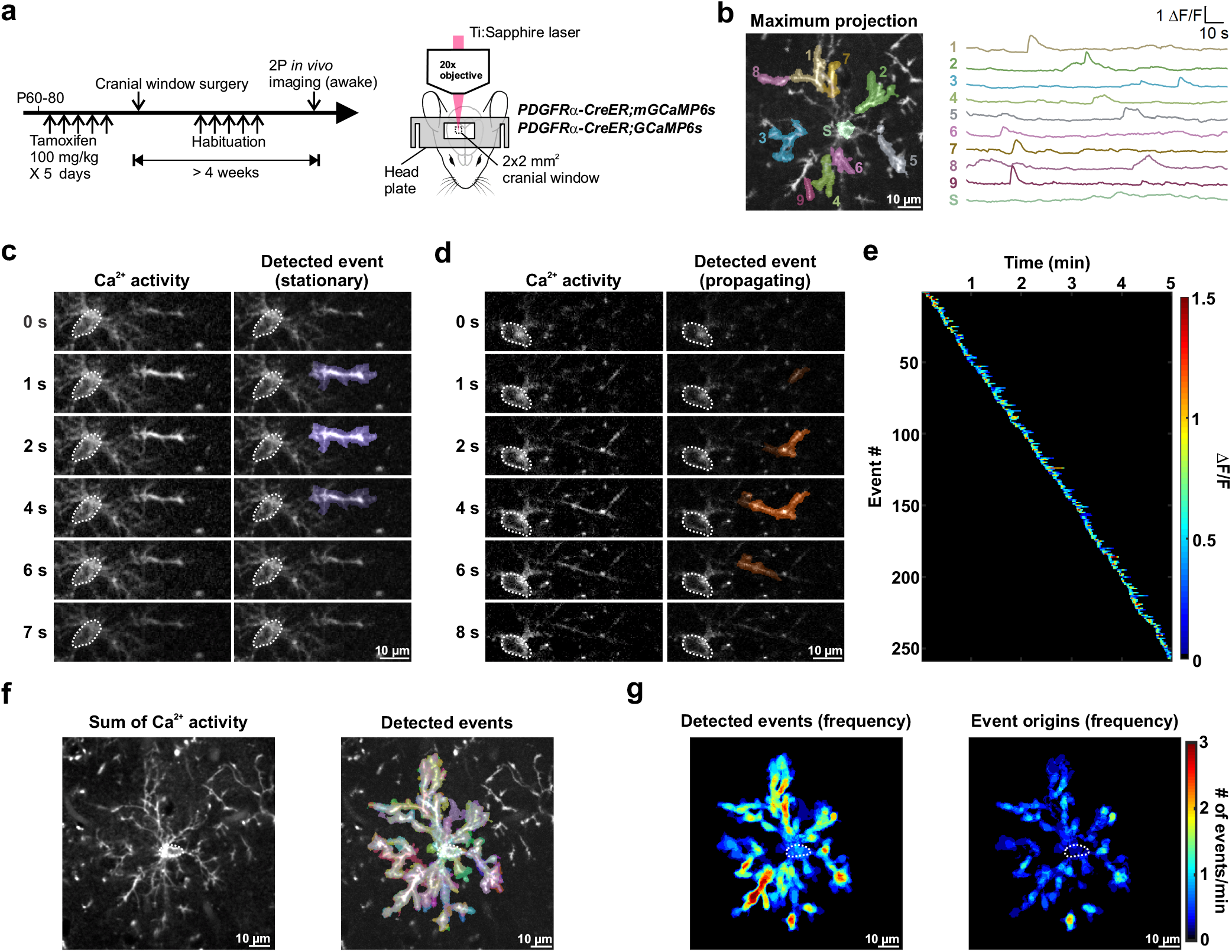
In vivo 2P imaging reveals the robust and dynamic Ca^2+^ activity in mouse cortical OPCs. **a**, Schematic illustrations of the research design. The expression of GCaMP6s in PDGFRα^+^ OPCs was induced between P60-80. OPC Ca^2+^ activity in the visual cortex of head-fixed, awake mice was observed through a chronic cranial window using 2P excitation from a Ti:Sapphire laser. See Methods for details. **b**, Example ΔF/F traces of OPC membrane Ca^2+^ events detected and randomly pseudocolored by AQuA software^31^. Example Ca^2+^ events were overlaid on the mGCaMP6s-expressing OPC visualized by projecting a total of 601 frames over 5 min onto a single image plane (Maximum projection). 1-9: process events. S: soma event. **c-d**, Frame-by-frame views of a stationary (c) and a propagating (d) OPC membrane Ca^2+^ event. Dotted white circles delineate the OPC cell body throughout. **e**, A heatmap showing all the Ca^2+^ events detected in an OPC over a 5-minute recording, sorted according to the time of the event onset. Note the absence of an overall increase of Ca^2+^ activity, which would suggest artificial photoactivation. **f**, Summation of all the membrane Ca^2+^ activities and events (randomly pseudocolored by AQuA) exhibited by an OPC during a 5-minute recording. Note how the events are wide-spread throughout OPC soma and processes. **g**, Heatmaps showing overall event frequency and event origins (defined as the location where the event reaches 20% of the amplitude) of the OPC in f.

Maps of summed activity superimposed on the cell revealed that Ca^2+^ events were not restricted to certain regions of the processes (see Fig. 1f); however, the amount of activity was not equally distributed, as regions of enhanced Ca^2+^ activity were apparent in maps of event frequency and event origin (the location when the event reached 20% of the peak ΔF/F) (Fig. 1g), suggesting that select regions are specialized to support this form of signaling. Propagating and non-propagating events occurred in the same processes, with propagating events comprising on average 39% of all Ca^2+^ transients (Extended data Fig. 3a). Some regions did not consistently produce propagating events (Extended Data Fig. 3b) and the distance that an event traveled did not correlate with its amplitude (Extended Data Fig. 3c), suggesting that propagation arises through active rather than passive processes. Moreover, events did not move in a consistent direction (i.e. to or from the soma) (Extended Data Fig. 3d), highlighting the variable nature of these events.

Activity in the processes of OPCs could trigger somatic Ca^2+^ events, integrating levels of activity to alter gene expression and cell behavior^32,33^. Indeed, less than 10% of all events occurred within the soma (GCaMP6s: 8.1 ± 1.0%; mGCaMP6s: 6.8 ± 1.6%; *n* = 6 each; p < 0.0001; Student’s t-test) (Extended Data Fig. 2b). If somatic events arise as a result of activity in the processes, there should be a temporal relationship between activity in these domains. To assess whether process and somatic events are correlated, we aligned all Ca^2+^ transients that occurred in a 20 s period in the processes relative to the onset of somatic events. Averages of activity in processes (Extended Data Fig. 3e, *solid lines*) indicated that somatic events were not preceded or followed by a consistent Ca^2+^ increase or pattern of Ca^2+^ transients in the processes. These results suggest that somatic Ca^2+^ increases arise from independent signaling events, rather than through an integration of process activity.

### OPC Ca^2+^ activity is suppressed during anesthesia

OPCs express a variety of neurotransmitter receptors that could influence Ca^2+^ dynamics during different brain states^17^. Volatile anesthetics such as isoflurane profoundly alter brain activity by suppressing neurotransmission and astrocyte Ca^2+^ signaling^34,35^. To assess whether OPC Ca^2+^ activity is also affected by changes in brain state, we performed *in vivo* 2P Ca^2+^ imaging and compared OPC activity before, during and after isoflurane-induced anesthesia. Anesthesia dramatically reduced the frequency of Ca^2+^ events by 72%, while the event area was reduced by ~50% (Fig. 2a, b). However, neither the amplitude nor the duration of Ca^2+^ transients were affected by anesthesia, suggesting that mechanisms responsible for internal Ca^2+^ regulation remain functional and that isoflurane may reduce the responsiveness of OPCs to external stimuli. Normal Ca^2+^activity patterns were still suppressed one hour after ceasing anesthesia, consistent with the lingering effects of general anesthetics on neuronal and astrocyte activity^34^, but recovered fully within 24 hours (Fig. 2b). These results indicate that OPC Ca^2+^ signaling is highly sensitive to anesthetics and correlated with the overall level of brain activity.

**Figure 2.**
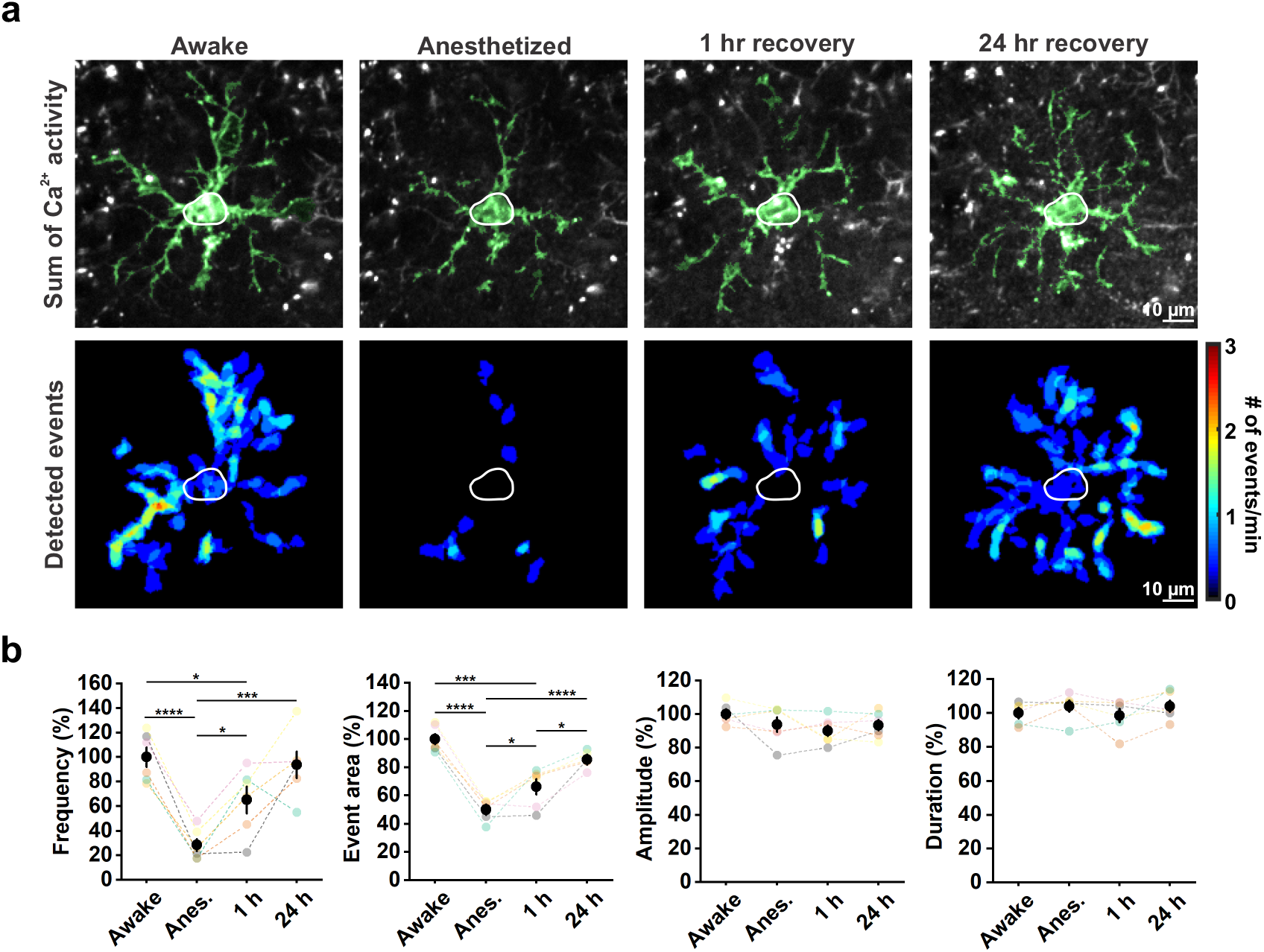
OPC Ca^2+^ activity is significantly suppressed during anesthesia. **a**, Representative images of a mGCaMP6s-expressing OPC exhibiting decreased Ca^2+^ activity during general anesthesia. The OPC morphology was visualized by summing a total of 361 frames (3 minutes) (Sum of Ca^2+^ activity) and highlighted in green. Heatmaps show Ca^2+^ event frequency (Detected events). **b**, Quantification of a. Both the frequency and event area were suppressed during general anesthesia while amplitude and duration of the events remained the same. *n* = 6 mice (color-coded). One-Way Repeated Measure ANOVA. Post-hoc: Tukey. *: p < 0.05; **: p < 0.01; ***: p < 0.001; ****: p < 0.0001 throughout.

### Enforced locomotion, but not light stimulation, stimulates OPC Ca^2+^ activity in the visual cortex

OPCs throughout the CNS express Ca^2+^-permeable ion channels and receive direct synaptic input from glutamatergic and GABAergic neurons^17–19^, raising the possibility that OPC Ca^2+^ transients are induced by neurotransmitter release from surrounding neurons. To address whether Ca^2+^ transients in OPCs arise through local neuronal activity, we activated neurons in visual cortex by exposing the contralateral eye to brief pulses of blue light in head-fixed, awake animals, while monitoring OPC Ca^2+^ activity in layer II of V1, where pyramidal neurons exhibit profound light-induced activity^36,37^ (Fig. 3a, b). Unexpectedly, Ca^2+^ transients in OPCs were unaffected by this intense visual stimulation (Fig. 3c-e), suggesting that local synaptic activity does not substantially contribute to the forms of OPC Ca^2+^ activity resolved by mGCaMP6s. However, on some trials light exposure was associated with enhanced Ca^2+^ activity, which occurred primarily when light stimulation startled the mice and induced ambulation, detected using an infrared (IR) camera mounted next to the stage during 2P imaging. To better define the influences of arousal/locomotion on OPC Ca^2+^ activity, we used a rotating, computer-controlled motorized platter to induce periodic enforced locomotion of the mice while imaging (Fig. 4a), a manipulation that reliably increases arousal and triggers widespread release of norepinephrine in the brain^36^. Using this paradigm, we observed that OPC Ca^2+^ activity was consistently enhanced during bouts of enforced locomotion (Fig. 4b, c and Supplementary Video 3). Ca^2+^ levels often increased throughout OPC processes and soma within seconds in response to this stimulation (Fig. 4d), resulting in an increase in event area (Fig. 4e, f). This increase of Ca^2+^ activity lasted for ~15-20 seconds (Fig. 4f), suggesting that the increased arousal state of the animals was responsible for the increased of Ca^2+^ activity in OPCs^36^. The amplitude of events was not increased by enforced locomotion (Fig. 4e, f), suggesting that this behavioral manipulation primarily increases the probability of that a Ca^2+^ transient will occur. Together, these data indicate that intracellular Ca^2+^ levels in OPCs are sensitive to brain state.

**Figure 3.**
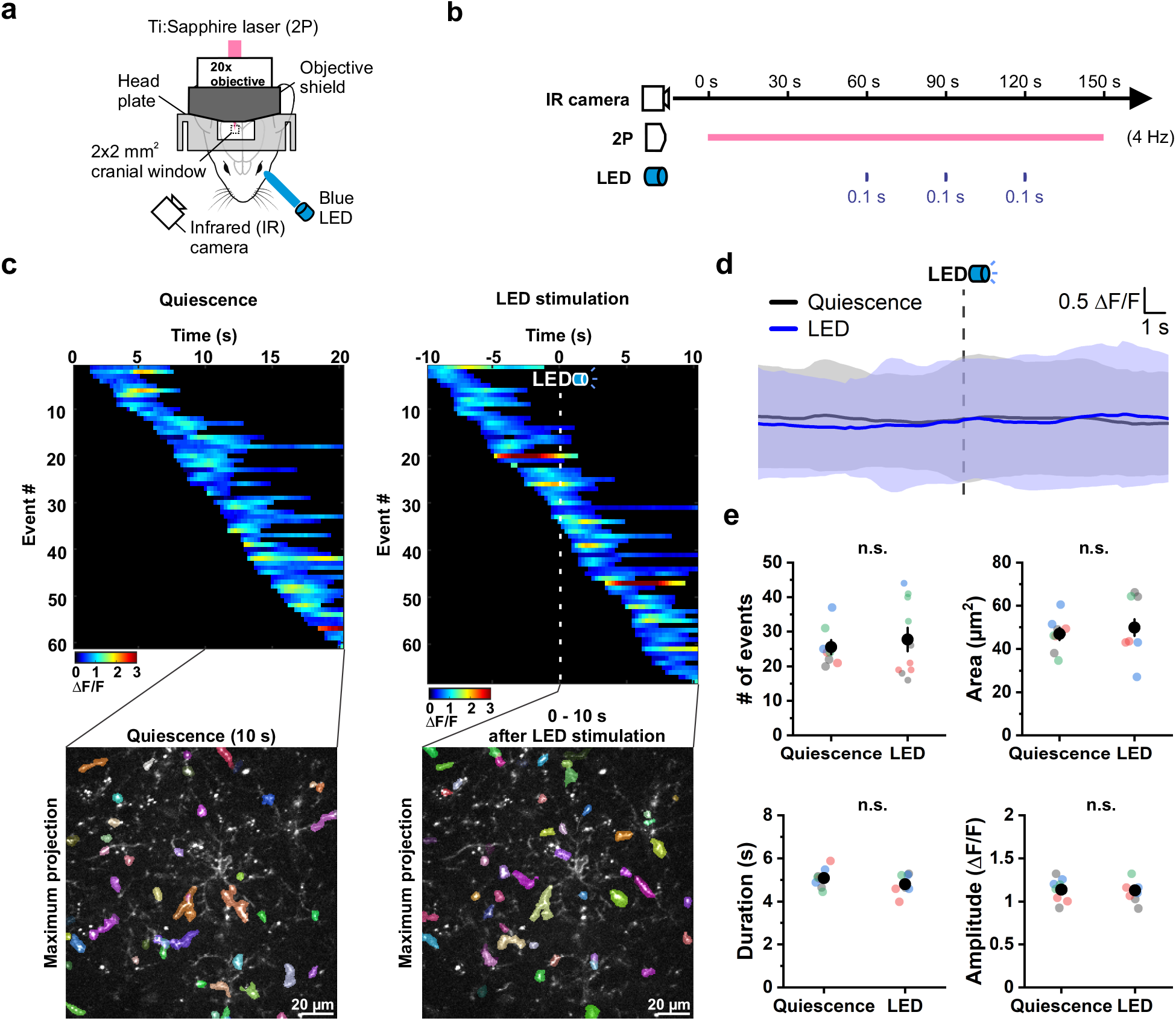
The excitation of the visual cortex by light does not affect OPC Ca^2+^ *in vivo*. **a**, Schematic illustration of the experiment setup. A customized 3D-printed objective shield was used to prevent LED light from entering the objective. The bottom part of the objective shield is not depicted in the illustration in order to display the cranial window. See Methods for details. **b**, Schematic illustration of the experiment design. Baseline OPC Ca^2+^ activity was recorded for 60 seconds (s) followed by 3 brief LED stimulations 30 s apart that lasted for 0.1 s each. An infrared (IR) camera was on throughout the experiment to observe mouse behaviors during image acquisition. **c**, Representative heatmaps showing the ΔF/F value and duration of OPC Ca^2+^ events sorted according to the time of event onset. OPC Ca^2+^ events that occurred during 10 s of quiescence or 10 s after LED stimulation were overlaid onto a single frame (Maximum projection), respectively. **d**, Averaging the OPC Ca^2+^ activity during 20 s of quiescence (gray) and around LED stimulation (blue) suggests that LED stimulation does not influence OPC Ca^2+^ activity *in vivo*. Shaded areas represent standard deviation. *n* = 4 mice. **e**, Quantification of OPC Ca^2+^ event frequency, area, duration and amplitude during 10 s of quiescence and 10 s post LED stimulation. *n* = 8 randomly-selected quiescent periods and 10 LED trials in 4 mice (color-coded). n.s.: not significant.

**Figure 4.**
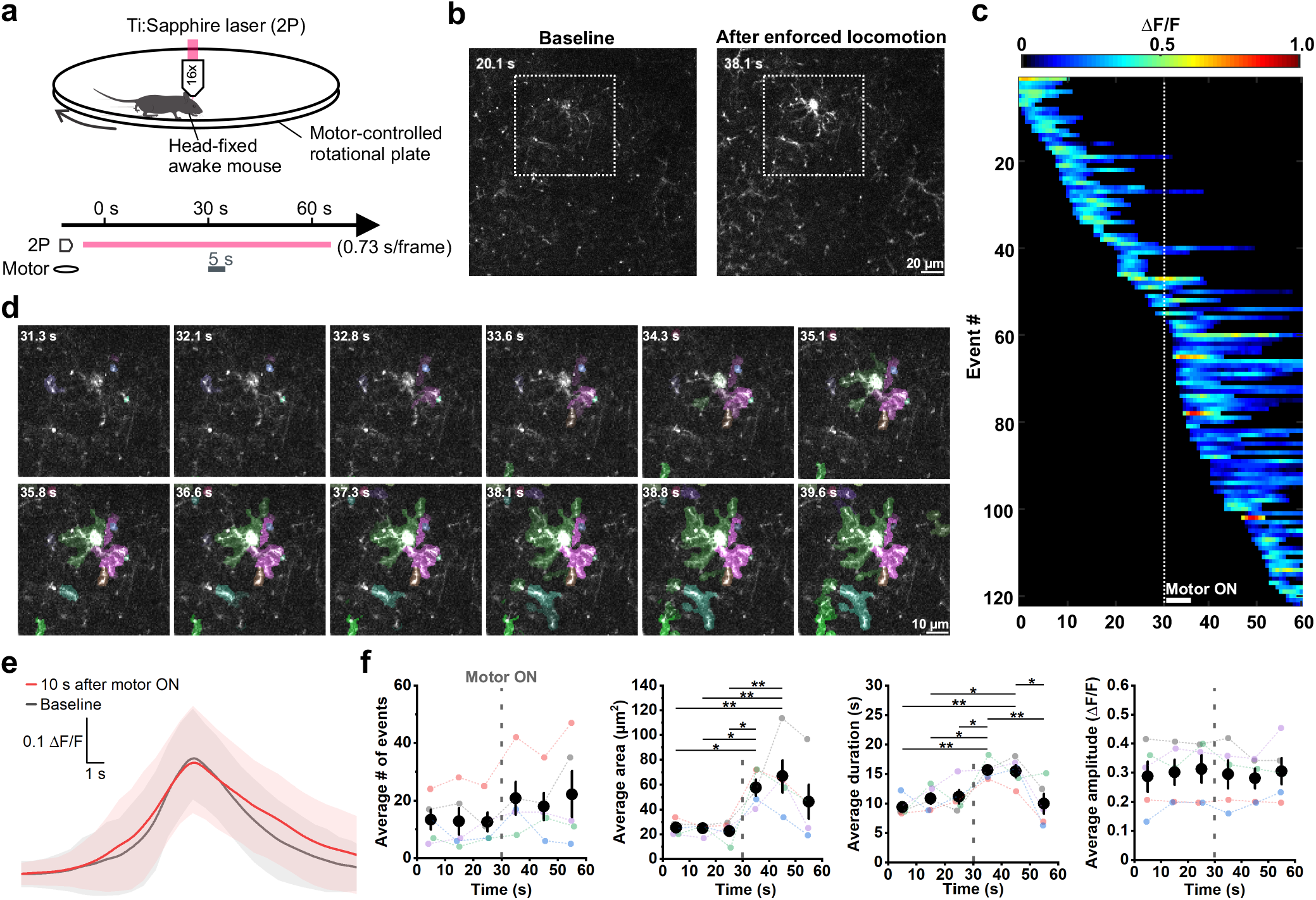
Enforced locomotion stimulates OPC Ca^2+^ activity in the mouse visual cortex. **a**, Schematic illustration of the experiment setup and design. During a 1 min recording window, enforced locomotion was triggered (Motor ON) at the 30^th^ s for 5 s. **b**, Representative 2P images showing OPC Ca^2+^ activity at baseline (20.1 s) and after enforced locomotion was triggered (38.1 s). **c**, A representative heatmap showing the OPC Ca^2+^ events sorted according to their time of onset during the 1 min recording. The dotted line indicates the time when the motor was turned on. **d**, Frame-by-frame views of the Ca^2+^ events (randomly pseudocolored) triggered by enforced locomotion in the white dotted square in b. Note the saturation of [Ca^2+^] increase in the processes and soma. **e**, Average ΔF/F traces of Ca^2+^ events that occurred during baseline (gray) and 10 s after the onset of enforced locomotion (red) aligned at their peaks. Shaded areas represent standard deviation. n = 114 baseline events and 131 enforced locomotion events in 5 mice. **f**, Quantification of the average number, area, amplitude and duration of the OPC Ca^2+^ events during the recording (binned by a 10-second interval). Gray dotted lines indicate the onset of enforced locomotion. One-Way Repeated Measure ANOVA. Post-hoc: Tukey. *n* = 5 mice (color-coded).

### Release of norepinephrine during arousal enhances OPC Ca^2+^ activity

The widespread enhancement of OPC activity during enforced locomotion suggests that these cells may be impacted by the global release of neuromodulators. Indeed, in quietly resting mice, the frequency of OPC Ca^2+^ events was strongly suppressed 20 minutes after i.p. administration of chlorprothixene (CPX) (Fig. 5), a sedative that inhibits a variety of neuromodulatory receptors. As transcriptional analyses suggest that serotonin, muscarinic, histamine and dopamine receptors are not highly expressed by OPCs^14,16,17^, we examined the involvement of noradrenergic signaling in OPCs by administering prazosin (Prz), an antagonist of α_1_ adrenergic receptor, which have been shown through *in vitro* pharmacology^38^, transcriptional profiling^14^ and reporter gene expression^39^ to be expressed by OPCs. Similar to CPX, *in vivo* administration of Prz also suppressed the frequency of Ca^2+^ transients in OPCs (see Fig. 5), suggesting noradrenergic signaling is a key modulator of OPC Ca^2+^ activity *in vivo*. Consistent with this hypothesis, OPC Ca^2+^ activity was also suppressed by *in vivo* administration of dexmedetomidine (Dex), an α_2_ adrenergic receptor agonist that inhibits norepinephrine release in the cortex. Similar to the effects of isoflurane-induced anesthesia (see Fig. 2), neither event amplitude nor duration were affected by these pharmacological manipulations, suggesting that adrenergic receptors primarily influence the probability of an event occurring, rather than the spatial and temporal profile of Ca^2+^ induced during an event. The size (area) of these events also trended lower when noradrenergic signaling was suppressed, although this decrease only reached statistical significance following administration of Dex. Together, these results show that OPC activity is modulated by noradrenergic signaling *in vivo*.

**Figure 5.**
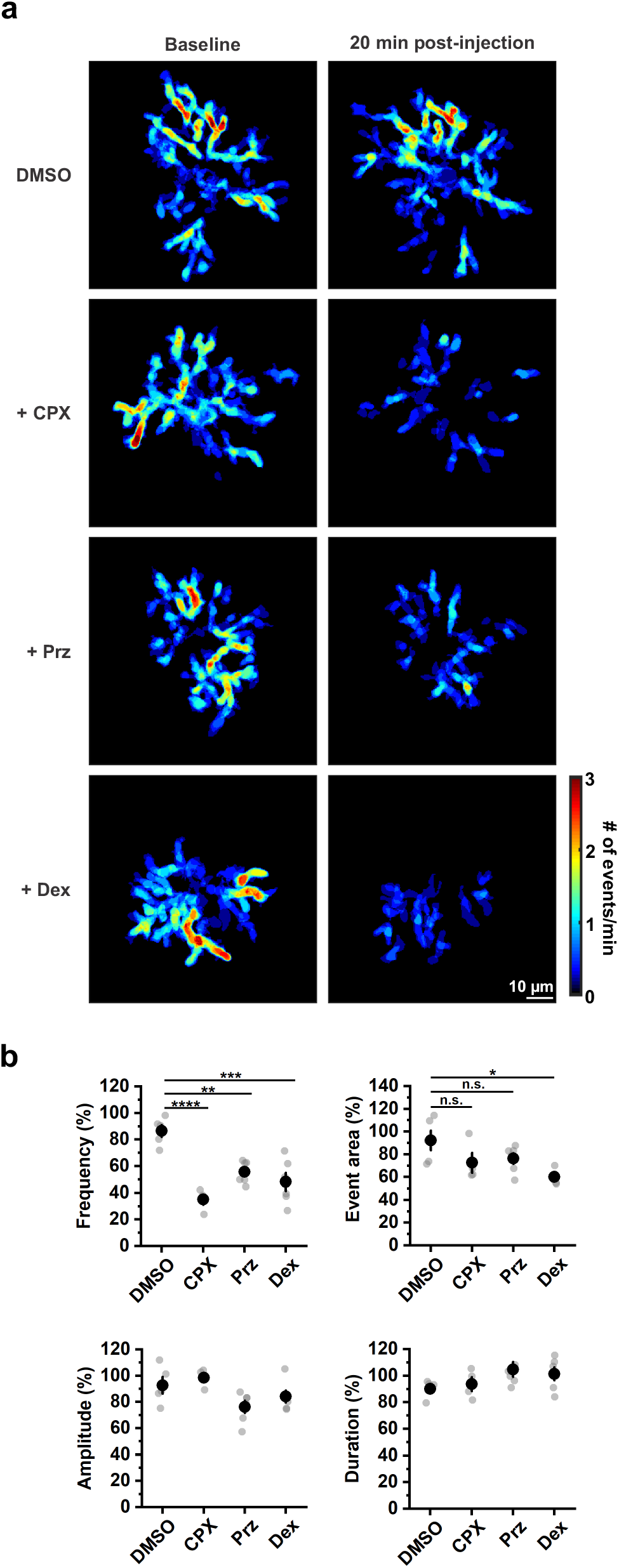
OPC Ca^2+^ activity is suppressed when the noradrenergic signaling is antagonized. **a**, Representative heatmaps showing the suppression of OPC Ca^2+^ activity by chlorprothixene (CPX, 5 mg/kg), prazosin (Prz, 3 mg/kg) and dexmedetomidine (Dex, 0.1 mg/kg), respectively. DMSO is used as the solvent control. **b**, Quantification of a. The frequency of the OPC Ca^2+^ events (% of baseline) was significantly suppressed by CPX, Prz and Dex compared to the control DMSO group. *n* = 5 mice for DMSO, 4 for CPX, 6 for Prz and 6 for Dex. One-Way ANOVA. Post-hoc: Tukey.

### OPCs are directly responsive to norepinephrine

Norepinephrine directly excites cortical neurons and enhances synaptic activity^40^, raising the possibility that the adrenoceptor-mediated enhancement of OPC Ca^2+^ activity observed *in vivo* may be indirect. To determine if norepinephrine has direct excitatory effects on OPCs, we measured the response of OPCs to adrenergic receptor agonists in acute cortical brain slices prepared from *PDGFRα-CreER;mGCaMP6s* mice. OPCs in these slices exhibited similar focal, but much less frequent and slightly longer duration Ca^2+^ transients in their processes compared to OPCs *in vivo* (Fig. 6a), differences that may reflect the lower temperature (room temperature vs. body temperature), and the preparation, in which connections with other brain areas are severed, notably between cortex and locus coeruleus where noradrenergic axons project from. To determine if these Ca^2+^ transients are dependent on synaptic activity, we superfused ACSF (artificial cerebral spinal fluid) containing tetrodotoxin (TTX, 1 μM), NBQX (10 μM), CPP (10 μM) and SR 95531 (20 μM) to block voltage-gated Na^+^ channels, AMPA receptors, NMDA receptors and GABA-A receptors, respectively, while monitoring OPC Ca^2+^ changes using 2P imaging. Remarkably, exposure to this broad combination of antagonists did not alter the frequency, area, or duration of spontaneous Ca^2+^ events in OPCs (Fig. 6b), suggesting that these events arise primarily from intrinsic processes, rather than engagement of neurotransmitter receptors during neuronal activity.

**Figure 6.**
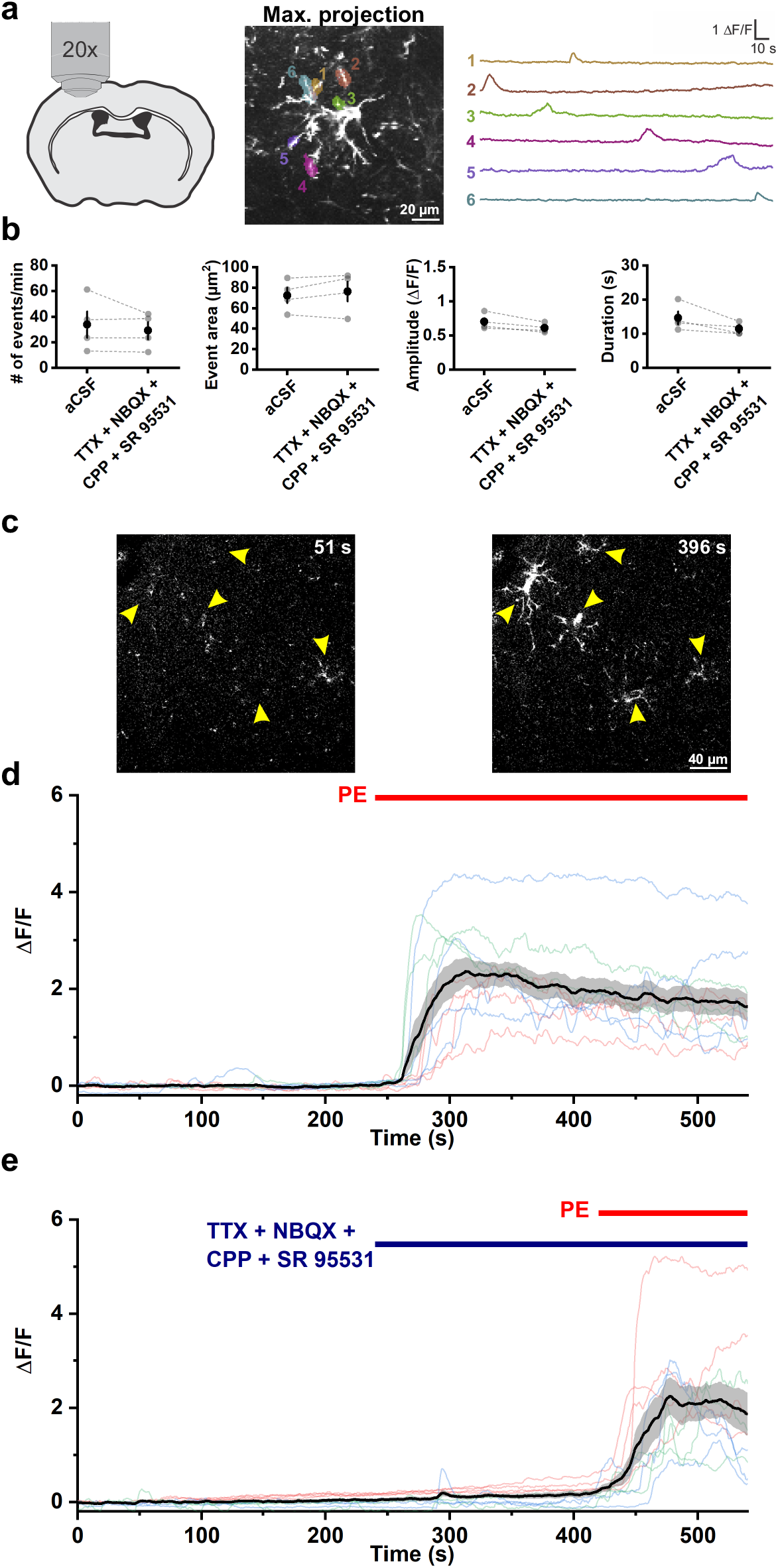
The OPC Ca^2+^ influx evoked by the α_1_ adrenergic receptor agonist is independent of synaptic activity *in vitro*. **a**, Example ΔF/F traces of OPC membrane Ca^2+^ transients observed in acute cortical slices in ACSF at RT. The morphology of the mGCaMP6s-expressing OPC was visualized by perfusing 10 μM of phenylephrine (PE) at the end of the recording (Also see in c). **b**, Comparing the average event frequency, area, amplitude and duration of the OPC membrane Ca^2+^ transients with and without the inhibitor cocktail containing: TTX (1 μM), NBQX (10 μM), CPP (10 μM) and SR 95531 (20 μM). *n* = 4 mice. Paired sample Student’s t-test. **c**, Representative images showing the evoked Ca^2+^ influx in the mGCaMP6s-expressing OPCs (yellow arrowheads) after the perfusion of PE (396 s) compared to ACSF only (51 s). **d**, Quantification of c. The black line and the shaded area represent mean ± SEM. *n* = 11 cells, 3 mice (color-coded). **e**, Quantification of the ΔF/F value from 10 OPCs in 3 mice (color-coded) when perfused with PE in the presence of the inhibitor cocktail.

To determine if direct activation of adrenergic receptors on OPCs is sufficient to alter Ca^2+^ activity, we exposed OPCs to the α_1_ adrenergic receptor agonist phenylephrine (PE, 10 μM). PE induced a dramatic increase in Ca^2+^ in OPCs in these acute brain slices (Fig. 6c, d and Supplementary Video 4). Similar to that observed with enforced locomotion (see Fig. 4), PE exposure led to Ca^2+^ elevations throughout the cell, making AQuA unsuitable for the analysis of discrete events. Therefore, we quantified the increase of intracellular Ca^2+^ (ΔF/F) based on the fluorescence change in each individual OPC as a region-of-interest (Fig. 6d and see Methods). To determine if synaptic release of glutamate and GABA contribute to the PE-induced OPC Ca^2+^ influx, PE was administered in the presence of the neuronal activity inhibitor cocktail (TTX, NBQX, CPP, SR 95531) (Fig. 6e). Consistent with the lack of change in OPC Ca^2+^ levels to local activity *in vivo* (see Fig. 3), neuronal activity inhibitors did not affect PE-induced OPC Ca^2+^ influx. Together, these results suggest that cortical OPCs express functional α_1_ adrenergic receptors that when activated trigger an increase in intracellular [Ca^2+^], providing an explanation for the increase in OPC Ca^2+^ activity observed during periods of enhanced arousal *in vivo*.

### α_1A_ adrenergic receptors mediate norepinephrine-associated OPC Ca^2+^ activity

The modulatory nature of the OPC response to norepinephrine and the slow time course of the Ca^2+^ increases induced by PE suggests that these effects are mediated by engagement of metabotropic receptors. To assess which alpha adrenoceptors could contribute to this activity, we examined the transcriptional profile of mouse cortical OPCs^14,16^. Among the nine different subtypes of adrenergic receptors, transcripts for Gq-coupled α_1A_ adrenergic receptors (*Adra1a*) were most abundant in OPCs^14,16^. Single molecule fluorescence *in situ* hybridization (smFISH) in brain tissue harvested from wild-type mice revealed that *Adra1a* mRNA co-localized with *Pdgfra* mRNA (Extended Data Fig. 4a, *white arrowheads*), suggesting that OPCs express α_1A_ adrenergic receptors *in vivo*, in accordance with the presence of EGFP^+^ OPCs in transgenic α_1A_-AR-EGFP reporter mice^39^. However, transcriptional profiling^14^, *in situ* hybridization (see Extended Data Fig. 4a), and functional analyses41 indicate that α_1A_ adrenergic receptors are not exclusively expressed by OPCs. Therefore, to determine if α_1A_ adrenergic receptors in OPCs mediate their responsiveness to norepinephrine, we conditionally knocked out *Adra1a* in OPCs by crossing *Adra1a^fl/fl^* mice^41^ with *PDGFRα-CreER;Rosa26-lsl-mGCaMP6s* mice (OPC α1A cKO mice). In acute slices prepared from these mice after TAM administration (see Methods), PE no longer elicited a rise in Ca^2+^ in the presence of neuronal activity blockers (TTX, NBQX, CPP, SR 95531), although they still responded to the Ca^2+^ ionophore ionomycin (Fig. 7a, b). To assess whether loss of α_1A_ adrenergic receptors also altered resting and evoked Ca^2+^ activity exhibited by OPCs *in vivo*, we analyzed their responses in the visual cortex of these mice using 2P imaging. OPC α1A cKO mice that were in a quiet resting state exhibited ~50% fewer OPC Ca^2+^ events compared to controls (Ctrl, *Adra1a^WT/WT^*) (Fig. 7c, d), suggesting that OPCs are subject to modulation by norepinephrine in the resting state. As observed previously, the amplitude, area and duration of these events were not affected, providing further evidence that norepinephrine primarily increases the probability of an event occurring, rather than being directly coupled to the Ca^2+^ rise itself. Deletion of *Adra1a* from OPCs prevented enhancement of OPC Ca^2+^ activity during enforced locomotion (Fig. 7e, f), indicating that activation of OPC α_1A_ adrenergic receptors are responsible for enhancing OPC Ca^2+^ activity during periods of arousal.

**Figure 7.**
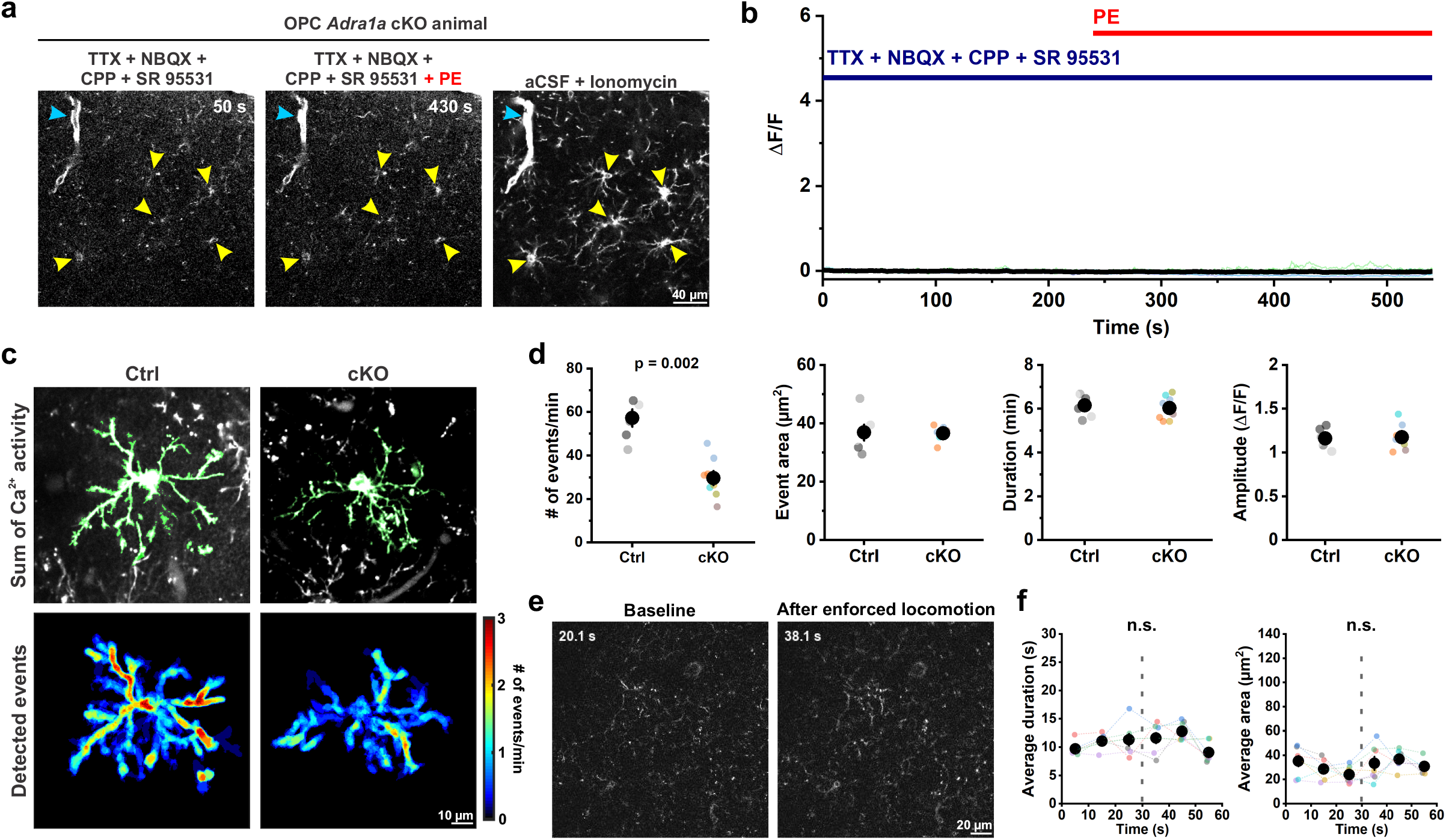
The removal of *Adra1a* from OPCs eliminates norepinephrine-mediated Ca^2+^ signaling. **a**, Representative images showing PE failed to evoke Ca^2+^ influx in the mGCaMP6s-expressing OPCs (yellow arrowheads) in acute cortical slices from *PDGFRα-CreER;Rosa26-lsl-mGCaMP6s;Adra1a^cKO/cKO^* animals (cKO). Note that PDGFRα^+^ perivascular fibroblasts (cyan arrowheads) were stimulated by PE but cKO OPCs were not. Ionomycin was applied at the end of recording to identify the mGCaMP6s-expressing OPCs. **b**, Quantification of a. The black line and the shaded area represent mean ± SEM. *n* = 18 cells, 3 mice (color-coded). **c**, Representative images and heatmaps showing cKO OPCs exhibited less frequent membrane Ca^2+^ transients than control (Ctrl, *PDGFRα-CreER;Rosa26-lsl-mGCaMP6s;Adra1a^wt/wt^*) animals. **d**, Quantification of c. n = 6/6 for Ctrl; 8/6 for cKO. Animals are color-coded. Student’s t-test. **e**, Representative images showing OPC Ca^2+^ activity in the cKO mice was not increased after enforced locomotion. **f**, Quantification of e. Gray dotted lines indicate the onset of enforced locomotion. *n* = 7 mice (color-coded). One-Way Repeated Measure ANOVA. Post-hoc: Tukey. n.s.: not significant.

### OPC Ca^2+^ activity is rapidly down regulated as they differentiate into myelinating oligodendrocytes

Examination of mRNA expression obtained from bulk RNAseq analysis of oligodendrocyte lineage cells (Brain RNAseq)^14^, revealed that *Adra1a* expression decreases as these cells began to differentiate, and becomes very low in mature oligodendrocytes, suggesting that within the oligodendrocyte lineage, OPCs may be particularly sensitive to modulation by norepinephrine. Consistent with this hypothesis, *in situ* hybridization in cortical brain sections revealed that *Adra1a* mRNA was less abundant in premyelinating oligodendrocytes (Extended Data Fig. 4b), identified by their expression of *lncOL1*, a long noncoding RNA specifically expressed at this stage of oligodendrocyte maturation^42^. To determine if noradrenergic modulation of Ca^2+^ activity declines with oligodendrocyte lineage progression, we performed longitudinal 2P Ca^2+^ imaging *in vivo* to monitor the Ca^2+^ activity of individual OPCs every other day for at least 28 days. Since OPCs continue to differentiate into myelinating oligodendrocytes in the adult cortex, we predicted that some tracked cells would begin this transition during the imaging period. Of the nine OPCs imaged from seven animals, four OPCs exhibited persistent Ca^2+^ activity throughout the imaging session, characterized by frequent localized increases in Ca^2+^ within their processes and soma (Fig. 8a, b, *green cell*; Fig 8d, *green lines*). Four other OPCs exhibited a precipitous decline in Ca^2+^ events (frequency, amplitude and duration) that began at different times during the imaging period (Fig. 8a, c, *red cell*; Fig. 8d, *red lines*), consistent with randomly timed lineage progression. The Ca^2+^ activity of one additional OPC also declined, but was followed by a dramatic increase of fluorescence, somatic swelling and process fragmentation, consistent with cell death (Extended Data Fig. 5; Fig. 8d, *gray line*).

**Figure 8.**
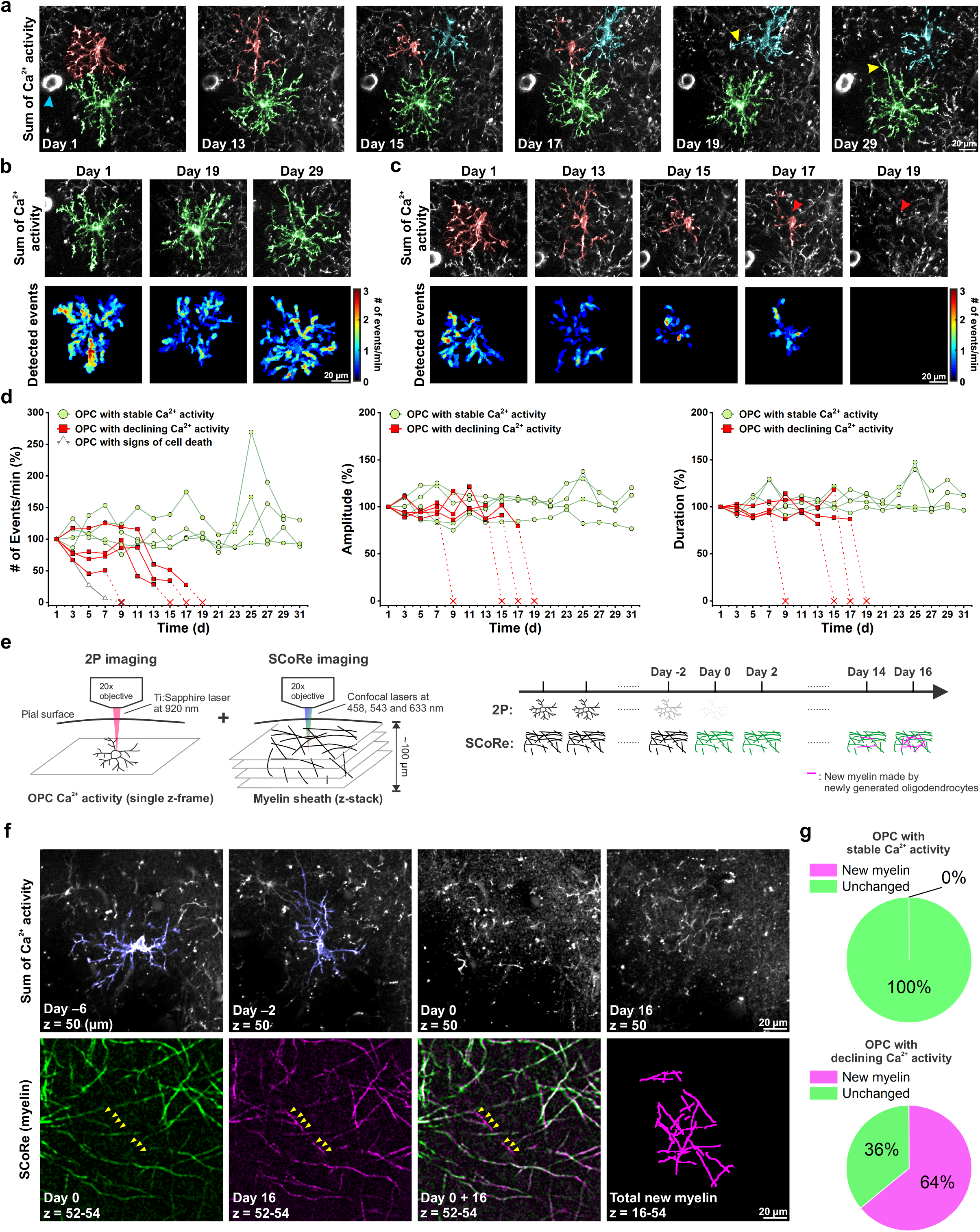
OPC Ca^2+^ activity declines gradually before differentiating into myelinating oligodendrocytes. **a**, Representative images showing diverse OPC Ca^2+^ behaviors across days. Cyan arrowhead indicates PDGFRα^+^ perivascular fibroblasts, which were used as landmarks to register images across different time points. The OPC highlighted in green persisted throughout the recording. The OPC highlighted in red became undetectable at Day 19, where a nearby OPC (cyan) extended its processes (yellow arrowhead) to the space now left unoccupied. A similar phenomenon was also seen with the neighboring green OPC at Day 29 (yellow arrowhead). **b**, Heatmaps showing the Ca^2+^ activity of the green OPC on Day 1, 19 and 29. **c**, Heatmaps showing the Ca^2+^ activity of the red OPC on Day 1, 13, 15, 17 and 19. Note how the frequency gradually declined since Day 15 and disappeared completely on Day 19. The red arrowhead indicates the position of the cell body of the disappearing red OPC on Day 17. **d**, Quantification of the OPC event frequency, amplitude and duration over time (*n* = 9 cells from 6 mice). Note only the frequency of Ca^2+^ events declined before the disappearance of OPCs, indicated by red Xs, but the amplitude and duration did not. **e**, Schematic illustration of the dual longitudinal imaging experiment using both 2P and SCoRe microscopies. Individual OPC Ca^2+^ activity was detected by 2P lasers at 920 nm. Meanwhile, the local myelin pattern surrounding the OPC of interest was recorded in a z-stack of SCoRe images using visible lasers. Dual imaging was performed every 2 days to observe the changes in OPC Ca^2+^ behaviors and local myelin pattern. After the OPC of interest became undetectable (Day 0), the dual imaging continued at the same location for 14-16 days to monitor the changes in local myelin pattern. **f**, An example of an OPC (highlighted in blue) with declining activity, and the local myelin pattern surrounding its cell body on the day the OPC became undetectable (Day 0, green), and 16 days after the disappearance (Day 16, magenta). Yellow arrowheads indicate the new myelin found 16 days after the disappearance of the OPC. **g**, Quantification of f. *n* = 18 cells (7 stable and 11 declining) from 4 mice.

OPCs maintain discrete territories in the CNS through repulsive interactions, and loss of one cell through differentiation or death is accompanied by rapid invasion of the territory of the lost cell by neighboring OPCs^4^. If the decline in Ca^2+^ activity observed in in OPCs was due to cell loss due to death or differentiation, then neighboring OPCs should have begun to extend their processes and invade the territory previously occupied by the missing cell. Indeed, this behavior occurred consistently for cells with declining Ca^2+^ activity (Fig. 8a, *cyan and green cells, yellow arrowheads*), suggesting that these changes in activity reflect loss of OPCs through differentiation and death.

To test the hypothesis that Ca^2+^ signaling rapidly declines during oligodendrocyte lineage progression, we simultaneously tracked Ca^2+^ activity within individual OPCs and local formation of myelin sheaths, by pairing *in vivo* 2P Ca^2+^ imaging in mice with SCoRE (<u>s</u>pectral <u>co</u>nfocal <u>re</u>flectance) imaging^43^, which allows visualization of myelin sheaths by imaging reflectance patterns. Ca^2+^ activity of OPCs in layer I of V1 was recorded through 2P imaging every other day within a single imaging plane. At the end of each session a z-stack of local myelin content was collected using SCoRE microscopy within the cell volume (50 μm above and below the 2P Ca^2+^ imaging plane) (Fig. 8e). Layer I OPCs were chosen for this analysis, because myelin sheaths are predominantly oriented horizontal to the cortical surface in this layer and therefore perpendicular to the incident light, which produces optimal reflection needed for SCoRE imaging^43^. Once Ca^2+^ activity in OPCs became undetectable (designated as Day 0), local myelin patterns continued to be visualized through SCoRE microscopy for another 14-16 days, the time that newly differentiated oligodendrocytes require to produce myelin sheaths^3,44^. For OPCs that exhibited stable Ca^2+^ activity (7/18 cells), no change in myelin patterns within these volumes were detected (Extended Data Fig. 6; Fig. 8g). In contrast, for 11 cells that exhibited a progressive decline in Ca^2+^ activity during the imaging period, new myelin sheaths appeared in 7 cells (64%) within the imaging volume 14-16 days later (Fig. 8f, g), supporting the hypothesis that oligodendrocyte maturation is associated with a dramatic decline in Ca^2+^ signaling. As OPCs underwent this transition, the frequency of Ca^2+^ events decreased, but the amplitude and duration were unaffected (see Fig. 8d), consistent with a decline in adrenergic modulation (see Fig. 5 and Fig. 7d). The four OPCs that did not produce new myelin sheaths may also have been differentiating, but failed to stably integrate, consistent with previous observations that the integration newly formed oligodendrocytes into cortical networks is inefficient^45^. Together, these results reveal that OPCs in the cerebral cortex are subject to brain state-dependent adrenergic modulation that rapidly declines as they mature into myelinating oligodendrocytes.

## Discussion

Neurotransmitters enable rapid signaling between neurons, enabling sensory processing, motor control and higher cognitive functions. Although glial cells are also responsive to neurotransmitters, we know comparatively little about the mechanisms used for neuron-glial communication or the consequences of this intercellular signaling. OPCs express a remarkable diversity of neurotransmitter receptors and their proliferation and differentiation are profoundly influenced by neural activity, suggesting that neurotransmitter signaling serves as a nexus to modify their homeostatic and regenerative behaviors. However, there have been no studies examining the physiological patterns of activity exhibited by OPCs in the intact brain to define which modes of communication are used to modify their properties during different behavioral states. Such physiological studies are crucial to define how their dynamics and ultimate fate are modulated by neurons. Here, we show using a new mouse line that enhances detection of near-membrane Ca^2+^ changes, that OPCs exhibit periodic increases in intracellular Ca^2+^ within their fine processes. This intrinsic activity was dramatically enhanced by norepinephrine, which activated α_1A_ adrenoceptors on OPCs during periods of enhanced arousal, providing a means to adapt OPC behavior to global changes in brain state.

OPCs are unique among glial cells in that they form direct synapses with neurons, serving exclusively as a postsynaptic partner. Prior studies have shown that direct activation of Ca^2+^-permeable ionotropic receptors^22,46–48^ or indirect activation of Ca^2+^ channels through depolarization^49,50^ can lead to cytosolic Ca^2+^ changes within OPC processes, and that Ca^2+^ signaling can modulate their migration, proliferation and differentiation^21,49,51,52^, as well as oligodendrocyte regeneration after CNS injury^53^. Thus, it is surprising that imaging with a membrane anchored sensor (GCaMP6s) did not reveal focal Ca^2+^ transients within OPC processes in the visual cortex when mice were exposed to intense light (see Fig. 3) or neuronal activity associated events in acute brain slices (see Fig. 6). Ca^2+^ flux through AMPA receptors occurs only when GluA2 subunits are absent, and expression of GluA2, Ca^2+^ permeable NMDA receptors and voltage gated Ca^2+^ channels vary among OPCs^54,55^ and functional expression of these channels has not been assessed in this region of the adult cerebral cortex. Moreover, detecting Ca^2+^ influx through AMPA receptors is challenging, because they exhibit rapid kinetics, producing highly focal Ca^2+^ transients^56^, and GCaMP must compete with endogenous buffers that may exhibit faster binding rates. Moreover, synaptic activation of AMPA receptors only leads to small depolarizations^18^, due to the high resting membrane K^+^ conductance exhibited by OPCs in the adult CNS^57^, which limits activation of voltage gated Ca^2+^ channels. It is also possible that these receptors are activated only under specific patterns of activity, such as burst firing, which may not have been induced while imaging.

In contrast to glutamate, GABA and acetylcholine, which are released at defined synaptic junctions, norepinephrine is released from varicosities along highly ramified axons that extend from neurons located in the locus coeruleus (LC)^58^. By utilizing a volume rather than direct synaptic mode of transmission, LC neurons can simultaneously modulate the activity of diverse targets within the vicinity of these projections. Previous studies indicate that glial cells are a key target of noradrenergic signaling^41,59,60^, as norepinephrine suppresses filopodial dynamics in microglia by activating β_2_-adrenergic receptors^59^, and enhances neurogenesis by activating β_3_-adrenergic receptors on radial glia in the hippocampus^60^, effects predicted to involve regulation of intracellular cAMP levels. Norepinephrine also has profound effects on astrocytes, triggering release of Ca^2+^ from intracellular stores by activating Gq-coupled α_1A_ adrenoceptors^41^. Engagement of astrocyte adrenergic receptors has been demonstrated *in vivo* in response to state transitions, such as sleep-wake^61^ and novel or unexpected experiences^36,62^, which result in widespread increases in intracellular Ca^2+^ throughout the astrocyte network, coinciding with widespread depolarization of cortical neurons. Exposure to norepinephrine enhances glyconenolysis in astrocytes^63^, raising the possibility that this pathway is used in a feed-forward manner to increase ATP production during periods of enhanced neuronal activity, anticipating increasing energetic demand for K^+^ and neurotransmitter uptake. Although activation of α_1A_ adrenoceptors on OPCs *in vitro* leads to cell-wide increases in Ca^2+^ similar to astrocytes (see Fig. 6), Norepinephrine release *in vivo* primarily enhances the magnitude of localized transients within their processes, amplifying intrinsic activity patterns (see Fig. 4). The distinct characteristics of OPC α_1A_ adrenoceptor signaling in the intact brain suggests that the concentration of norepinephrine that reaches these receptors *in vivo* is lower than can be achieved through exposure to exogenous agonists. Although *in vitro* studies suggested that oligodendrocytes continue to express α_1A_ adrenergic receptors and exhibit Ca^2+^ responses to norepinephrine^11,64^, transcriptional profiling indicates that these receptors are downregulated with development and our studies indicate that both intrinsic and norepinephrine-enhanced Ca^2+^ signaling decline rapidly with differentiation (see Fig. 8). Thus, OPCs appear to be uniquely specialized within the oligodendrocyte lineage to respond to norepinephrine.

As expected from the widespread activation of LC neurons during state transitions, OPCs throughout the imaging field exhibited similar activity enhancement during these behaviors (see Fig. 4), indicating that norepinephrine can exert a widespread modulatory effect on these progenitors. Assessments of transcriptional changes using translating ribosome affinity purification (TRAP) and mRNA microarray analysis^65^, indicate that OPCs exhibit circadian changes in gene expression, with transcription of genes associated with cell proliferation and growth increasing during sleep, when the brain norepinephrine is low, and genes associated with lineage progression during wakefulness, when the brain norepinephrine level is high^65^. Thus, norepinephrine may bias the behavior of OPCs, supporting restorative cell homeostasis during sleep, and oligodendrogenesis during wakefulness. Recent studies indicate that other neurotransmitter receptors, including Kappa opioid receptors^23^ and metabotropic M1 acetylcholine receptors^24^ expressed by OPCs can also influence their lineage progression, suggesting that these progenitors have the ability to sense the activity patterns of distinct subsets of neurons. Although it is not yet clear how they integrate these diverse signals, behavioral manipulations such as intense motor learning can mobilize these progenitors to differentiate and produce additional myelin^66^, a phenomenon termed adaptive myelination. Such training paradigms induce a heightened state of arousal and are associated with increased activation of the LC and cortical norepinephrine release. Expression of α_1A_ adrenoceptors by OPCs may allow these progenitors to sense these state changes and lower barriers for differentiation, enabling these cells to modify neural circuits in response to changes in life experience.

## Acknowledgements

We thank our friends and colleagues for their input and support throughout the study: T. Babola, Y. Wang and G. Yu all provided helpful suggestions for analyzing OPC Ca^2+^ activity. C. Call provided assistance in SCoRe microscopy. N. Ye and A.E. Bush helped with mouse husbandry. M. Pucak and A.E. Bush provided assistance with daily operation and maintenance of the microscopes essential to this study. We appreciate the generosity of the Scidraw community, especially Luigi Petrucco (doi.org/10.5281/zenodo.3925903, used in Fig. 1a), Ethan Tyler Lex Kravitz (doi.org/10.5281/zenodo.3925975, used in Fig. 3a and Extended Data Fig. 5d), and Agustin Carpaneto (doi.org/10.5281/zenodo.3926119, used in Fig. 5). This study was supported by grants from the Dr. Miriam and Sheldon G. Adelson Medical Research Foundation (AMRF) and the National Institute of Aging (AG072305).

## Author contributions

T.-Y.L. and D.E.B. designed the research and experiments. T.-Y.L. performed and analyzed all the experiments. P.H. contributed to the analysis of the longitudinal OPC Ca^2+^ activity. E.T.H. constructed the enforced locomotion rig. A.A. designed and generated the *Rosa26-lsl-cGCaMP6s* and *Rosa26-lsl-mGCaMP6s* transgenic mouse lines. T.-Y.L., P.H. and D.E.B. co-wrote the manuscript.

## Materials & Correspondence

To inquire about the materials used in this study, please contact D.E. Bergles.

## Extended Data Figure Legends

**Extended Data Figure 1.**
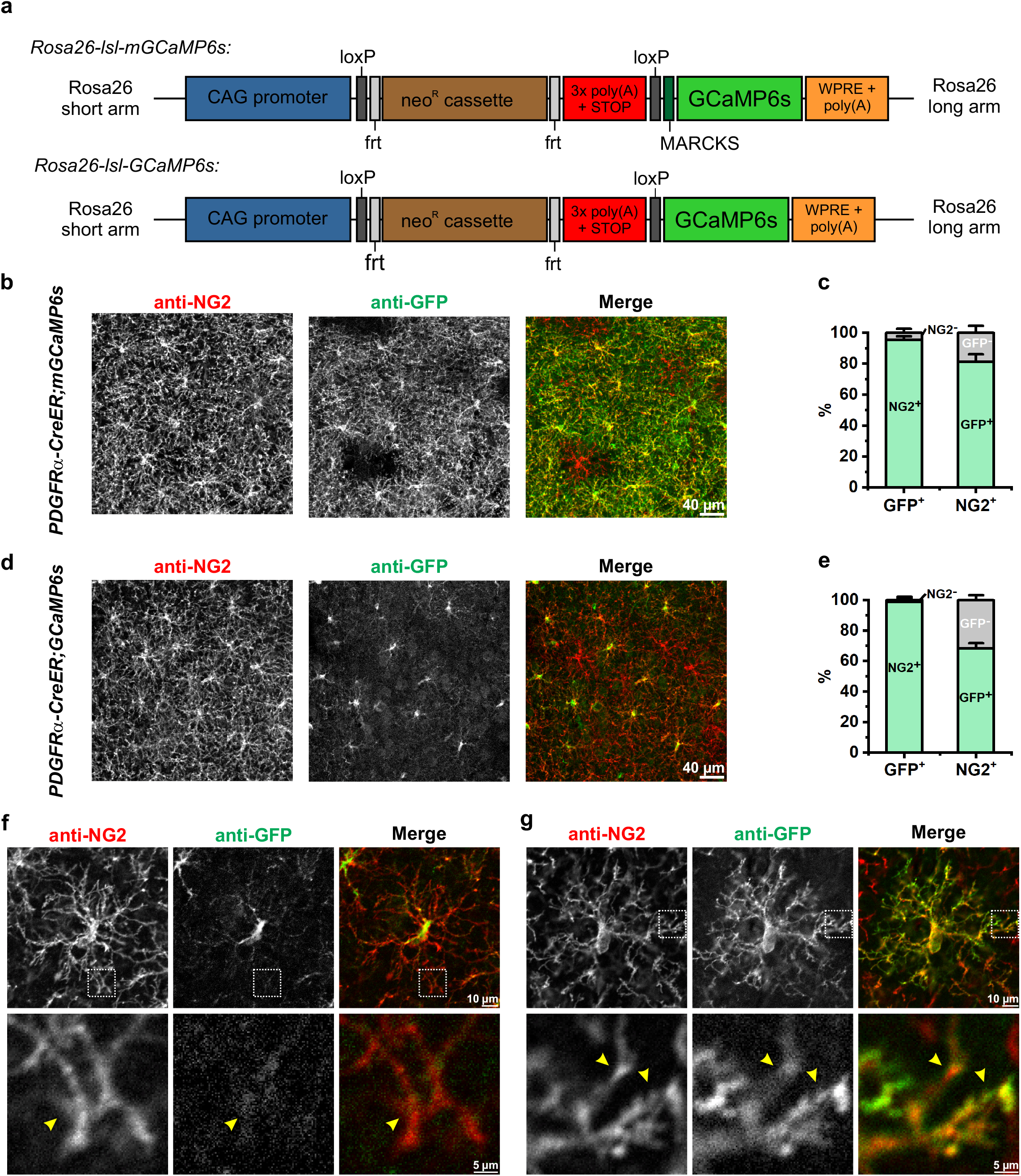
Expressing membrane-tethering GCaMP6s (mGCaMP6s) in OPCs using the *Rosa26-lsl-mGCaMP6s* knockin transgenic mice. **a**, The design of the *Rosa26-lsl-mGCaMP6s* and *Rosa26-lsl-GCaMP6s* knockin transgenic mice. MARCKS: the N-terminal myristoylation sequence of <u>m</u>yristoylated <u>a</u>lanine-<u>r</u>ich <u>C</u>-<u>k</u>inase <u>s</u>ubstrate. **b**, Representative confocal images showing the expression of mGCaMP6s (anti-GFP) in the cortical OPCs (anti-NG2) following immunohistochemistry 4 weeks after tamoxifen injection. **c**, Quantification of b (*n* = 3 mice). **d**, Representative confocal images showing the expression of cytosolic GCaMP6s in the cortical OPCs following immunohistochemistry 4 weeks after tamoxifen injection. **e**, Quantification of d (*n* = 3 mice). **f-g**, Representative confocal images of single mGCaMP6s-(f) and GCaMP6s-expressing (g) OPCs with mafnified views of their distal processes (yellow arrowheads) in the dotted squares, respectively. Note the lack of cytosolic GCaMP6s expression in the processes.

**Extended Data Figure 2.**
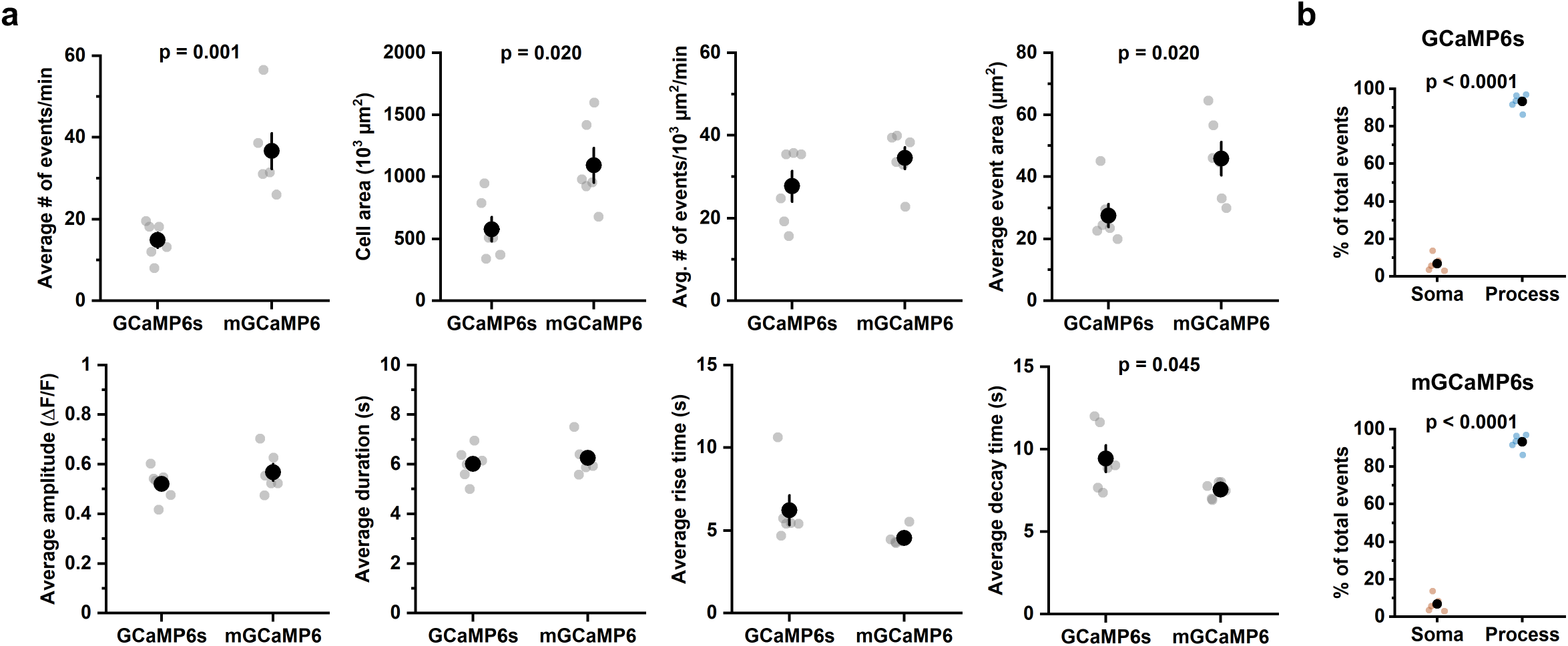
Basic properties of OPC Ca^2+^ events detected by GCaMP6s and mGCaMP6s, respectively. **a**, Box plots comparing the average event frequency (events/min), cell area, frequency normalized to cell area, event area, amplitude, duration (the time between 50% onset time point – 50% offset time point), rise time (onset duration from 10% to 90% of the peak amplitude) and decay time (offset duration from 90% to 10%) between GCaMP6s-expressing and mGCaMp6s-expressing OPCs. *n* = 6 mice each. Student’s t-test. **b**, Quantification of the event origins from GCaMP6s-expressing and mGCaMP6s-expressing OPCs (*n* = 6 mice each). Black filled circles and whiskers represent mean ± SEM. Student’s t-test.

**Extended Data Figure 3.**
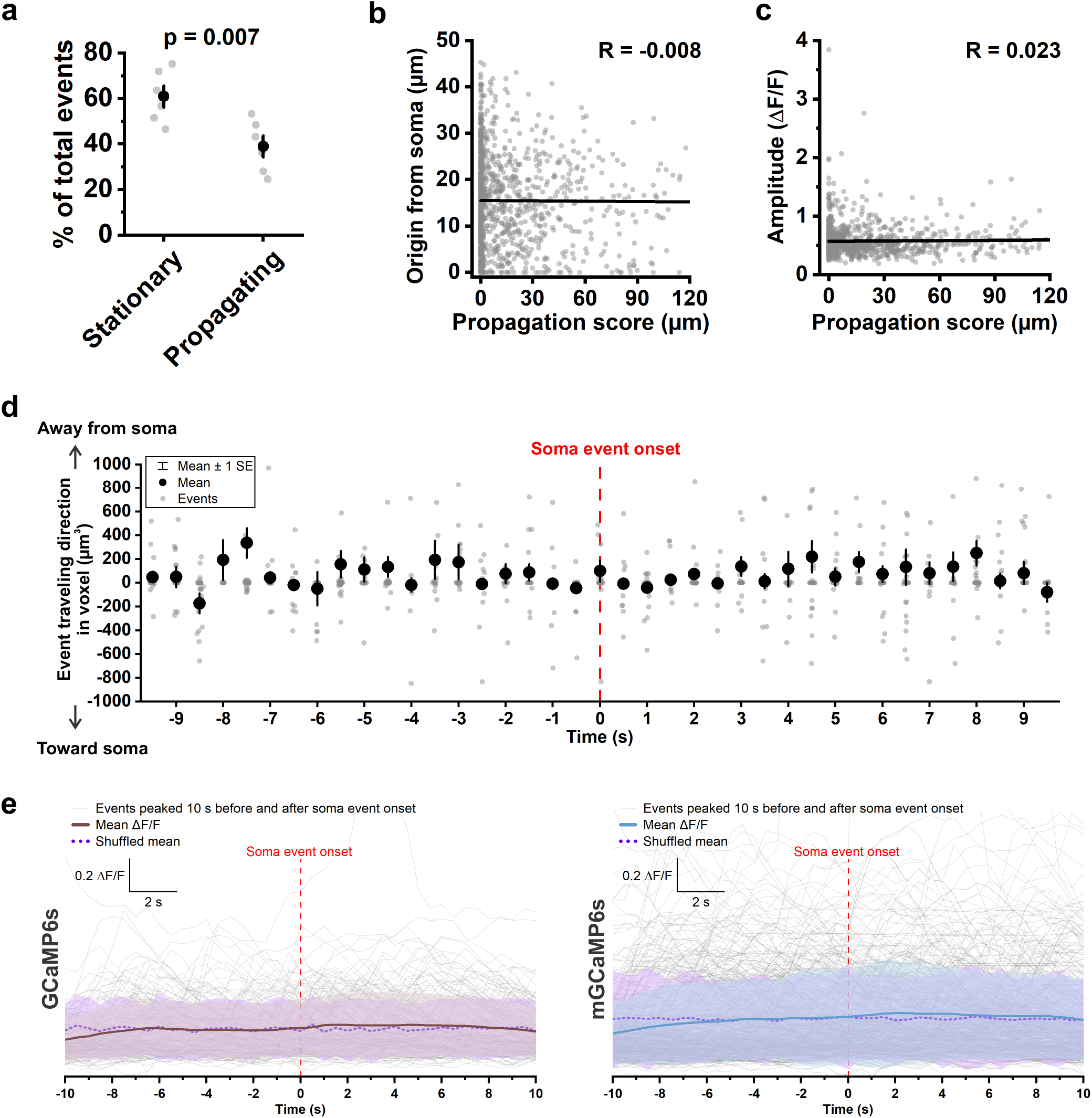
Membrane OPC Ca^2+^ transients propagate independent of the location of event origin, event amplitude, and the Ca^2+^ activity in the soma. **a**, The average percentage of stationary (having an overall propagation score < 10 μm) and propagating (having an overall propagation score ≥ 10 μm) OPC membrane Ca^2+^ events (*n* = 6 mice). Student’s t-test. **b**, A plot of the distance between event origin and soma (Origin from soma) versus overall propagation score. R: Pearson’s r. *n* = 1,100 propagating events from 6 mice. **c**, Plotting event amplitude against event overall propagation score. R: Pearson’s r. *n* = 1,100 propagating events from 6 mice. **d**, Directions of event propagation 10 s before and after the onset of a soma event. Event travelling direction was determined by total voxels travelled away from soma minus total voxels travelled toward soma. *n* = 911 propagating events from 6 mice. **e**, ΔF/F traces of the process events (thin gray lines) peaked 10 s before and after soma event onset in GCaMP6s-expressing and mGCaMp6s-expressing OPCs, respectively. Mean ΔF/F (solid brown and blue lines) is the average ΔF/F of 165 events in the cGCaMP6s-expressing mice (6 cells from 6 mice), and 509 events in the mGCaMP6s-expressing mice (6 cells from 6 mice). Shuffled mean (dotted purple lines) is the average value after shuffling ΔF/F values of each event. Shaded areas indicate standard deviation.

**Extended Data Figure 4.**
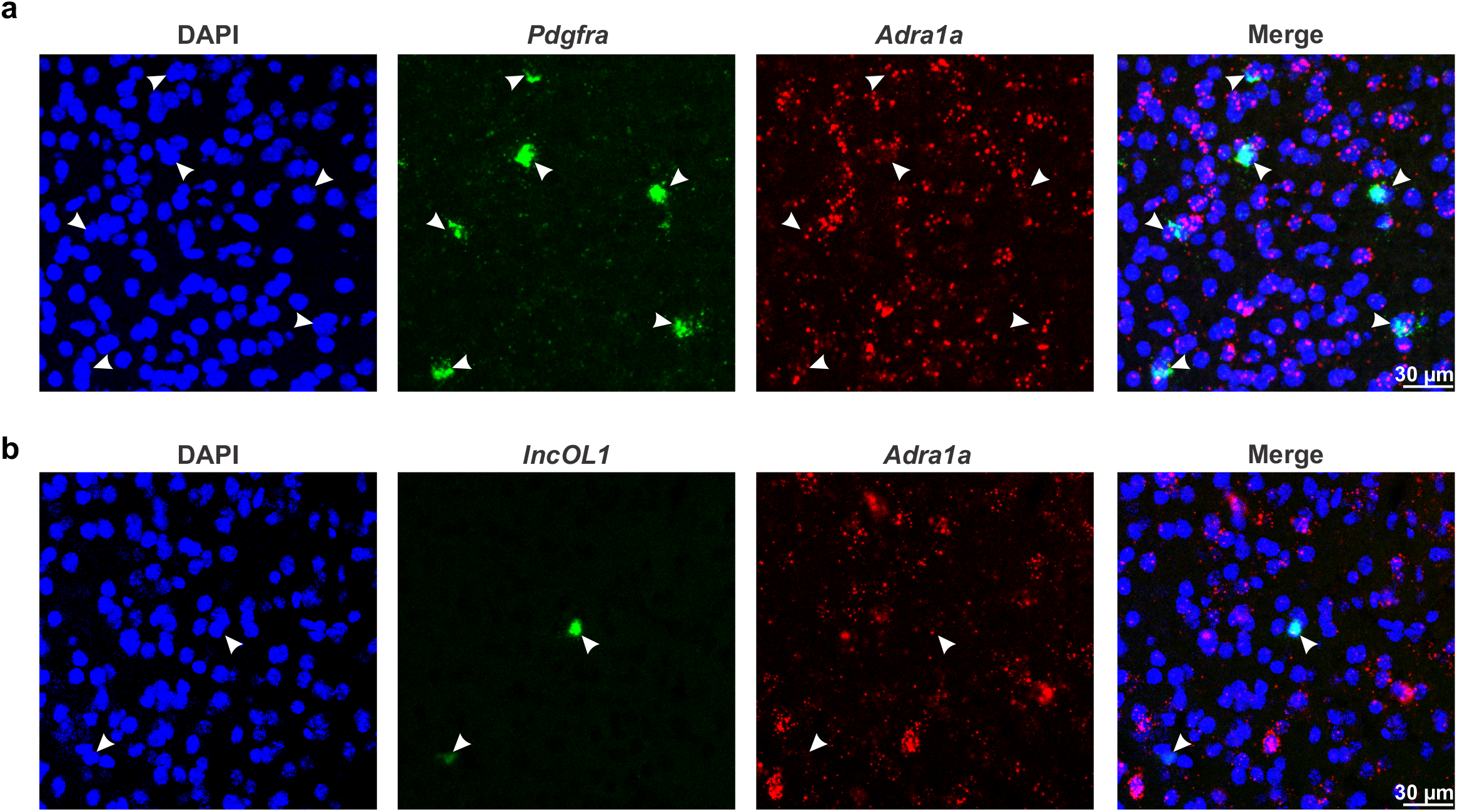
The mRNA of the α_1A_ adrenergic receptors are abundantly expressed in OPCs but not in pre-myelinating oligodendrocytes. **a**, Representative confocal images of an adult B6 visual cortex hybridized with probes recognizing the OPC marker, *Pdgfra* (green) and *Adra1a* (red) mRNA. DAPI (blue) stains cell nuclei. *Adra1a* mRNA is found around *Pdgfra*^+^ nuclei, suggesting that cortical OPCs express ADRA1A (*n* = 3 mice). **b**, Representative confocal images of an adult B6 visual cortex hybridized with probes recognizing the pre-myelinating oligodendrocyte marker, *lncOL1* (green), and *Adra1a* (red) mRNA. DAPI stains cell nuclei (*n* = 2 mice).

**Extended Data Figure 5.**
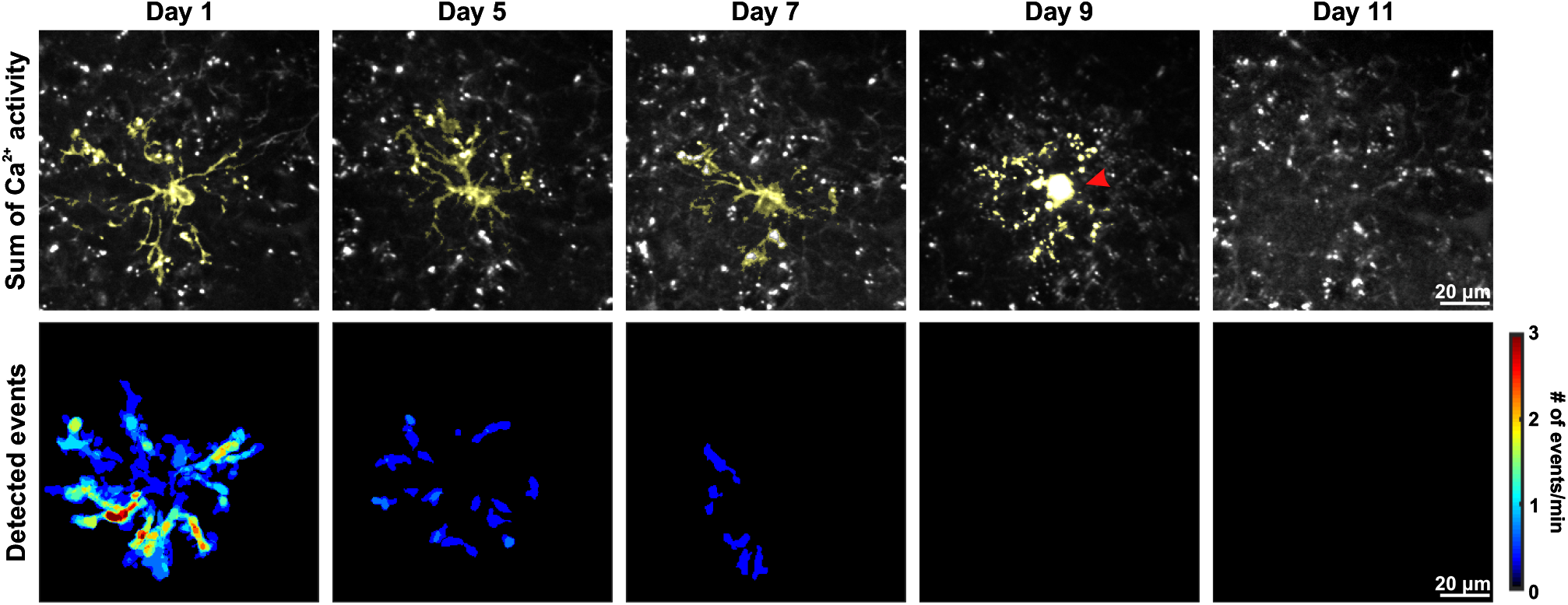
An example of a mGCaMP6s-expressing OPC undergoing cell death. The mGCaMP6s-expressing OPC is highlighted in yellow. Note the round-shaped and intensely bright cell body (red arrowhead) as well as the fragmented processes on Day 9. We could not identify any Ca^2+^ events in the fragmented processes.

**Extended Data Figure 6.**
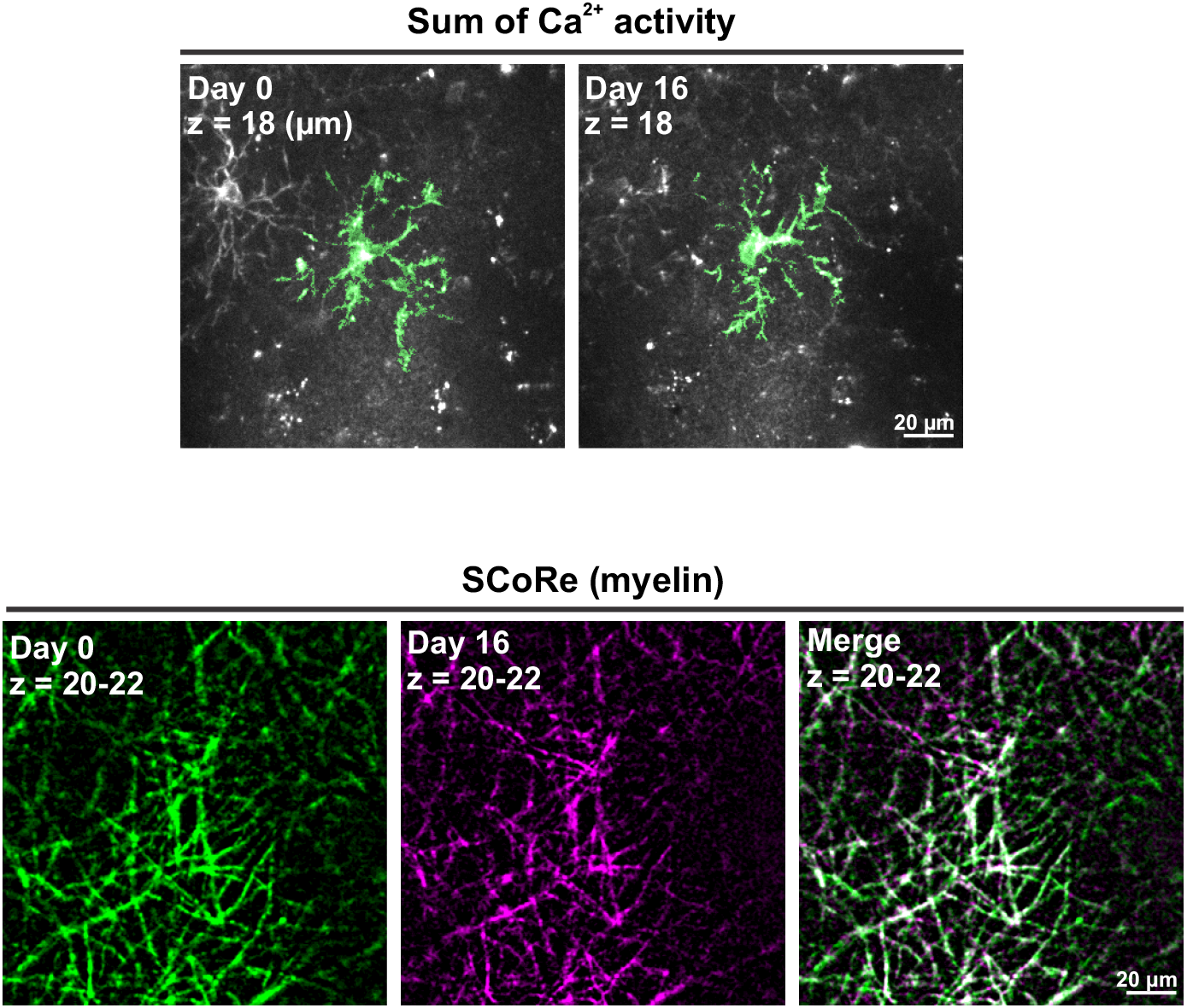
An example of local myelin profile unchanging around a stable OPC. The mGCaMP6s-expressing OPC (highlighted in green) was followed for 16 days and local myelin profile was recorded by SCoRE microscopy concurrently. Local myelin profile remained unchanged from Day 0 (green) to Day 16 (magenta).

**Extended Data Figure 7.**
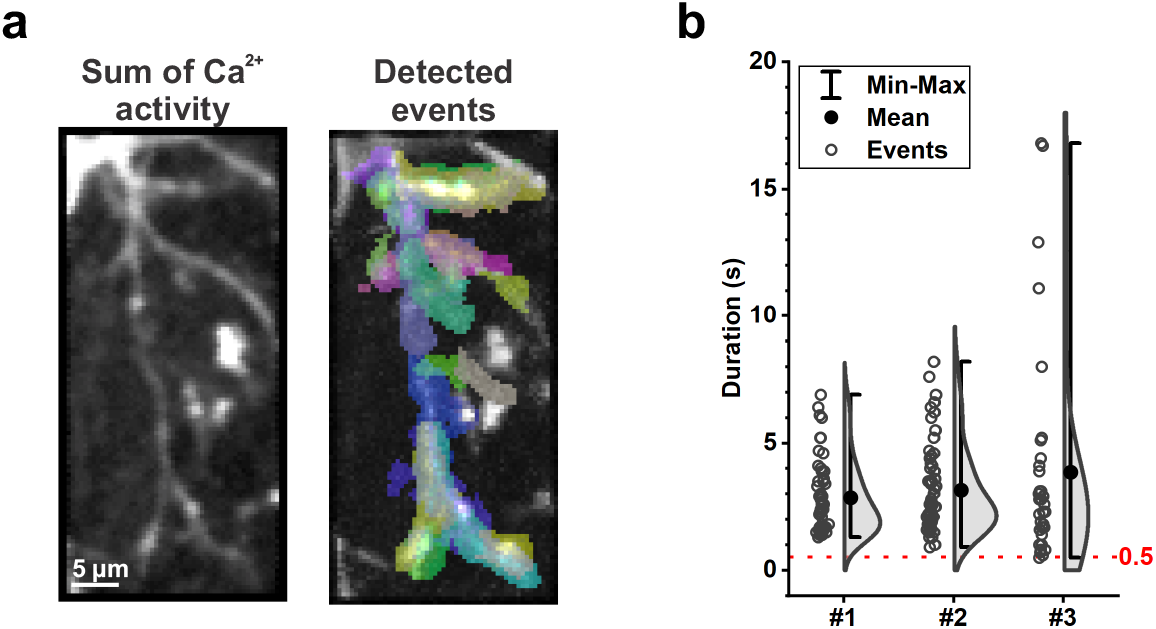
OPC Ca^2+^ events last a minimum of 0.5 seconds. **a**, Scan and analysis of Ca^2+^ events in OPC processes at 10 Hz (Cell body at north). **b**, Quantification of Ca^2+^ event duration, detected using 20 Hz (animal #1) or 10 Hz (animal #2 and #3) of 2P Ti:Sapphire lasers at 920 nm. Note that every event detected is at least equal to or longer than 0.5 s.

## Methods

### Animal care and use

Female and male adult mice were used for experiments and randomly assigned to experimental groups. No noticeable sex-specific behavioral or physiological phenotypes were observed throughout the research. All mice were healthy and none were excluded from the analysis. Mice were maintained on a 12-hour light/dark cycle, housed in groups no larger than five, and received food and water ad libitum. All animal experiments were conducted in accordance with the National Institute of Health Guide for the Care and Use of Laboratory Animals and protocols approved by the Animal Care and Use Committee at Johns Hopkins University.

### Transgenic animals

Conditional knockin *Rosa26-lsl-GCaMP6s* and *Rosa26-lsl-mGCaMP6s* mouse lines were generated by inserting a conditional allele into the Rosa26 locus (see Extended Data Fig. 1). The *PDGFRα-CreER* BAC transgenic mouse line was described in Kang et al^1^. The *Adra1a^fl/fl^* transgenic mouse line was described in Ye et al^41^.

### Tamoxifen preparation and administration

To induce GCaMP6s expression in *PDGFRα-CreER;GCaMP6s* and *PDGFRα-CreER;mGCaMP6s* transgenic mice, TAM (Sigma-Aldrich, T5648) was freshly prepared on the first day of the injection at 10 mg/mL in sunflower seed oil (Sigma-Aldrich, S5007) through intermittent sonication at room temperature (RT). Adult (> 8 weeks) mice were injected intraperitoneally (i.p.) with a dosage of 100 mg/kg body weight (b.w.) for five consecutive days, once per day. Every injection was at least 20 hours (hrs) apart. The remaining tamoxifen solution was stored at 4°C in the dark for a maximum of 5 days. All experiments were performed at least 2 weeks after the last tamoxifen injection.

### Immunohistochemistry

Mice were deeply anesthetized with an i.p. injection of pentobarbital (100 mg/kg b.w.) and perfused transcardially with 20 mL of RT 0.1 M phosphate buffered saline (1x PBS) first, and then with 20 mL of freshly prepared, ice-cold 4% paraformaldehyde (PFA, Electron Microscopy Sciences, #19210) in 1x PBS (pH7.4). After carefully removing from the skull, brains were post-fixed in 4% PFA/PBS at 4°C in the dark for 4 hrs, and then cryoprotected in 30% sucrose/0.1 % sodium azide in 1x PBS at 4°C in the dark for at least 48 hrs. Brains were then embedded in Tissue-Tek O.C.T Compound (Sakura Finetek, #4583) and cyrosectioned at −20°C using a Thermo Scientific Microm HM 550 at the thickness of 35 μm. Free-floating coronal brain sections were collected and rinsed briefly in PBS and then permeabilized with 0.5% Triton X-100 in 1x PBS for 10 min at RT for immunostaining. To prevent non-specific binding of antibodies, brain sections were incubated in blocking buffer (10% normal donkey serum, Jackson ImmunoResearch, #017-000-121, and 0.3% Triton X-100 in 1x PBS) for 1 hr at RT, followed by primary antibody incubation at RT overnight. After washing in 1x PBS 3 times, 10 min per wash, brain sections were incubated with secondary antibodies for 2 hours at RT before another wash in 1x PBS as described above. Both primary and secondary antibodies were diluted in the blocking buffer. Sections were then mounted on slides with Aqua-Poly/Mount (Polysciences, #18606). Images were acquired using a Zeiss LSM 800 confocal microscope with Zeiss 10x/0.45 Plan-Apochromat and Zeiss 20x/0.8 Plan-Apochromat objectives and analyzed using ImageJ (National Institute of Health). Primary antibodies used in this study were guinea pig anti-NG2 (Bergles lab, 1: 5000)^67^ and chicken anti-GFP (Aves, #GFP-1020, 1:4000). Secondary antibodies used in this study were purchased from Jackson ImmunoResearch and used at 1:1000: Cy3 donkey anti-guinea pig IgG (#706-165-148) and Alexa Fluor 488 donkey anti-chicken IgG (#703-546-155).

### Single molecule fluorescent in situ hybridization

B6 wild-type adult mice were deeply anesthetized with an i.p. injection of pentobarbital (100 mg/kg b.w.) and perfused transcardially with 20 mL of RT 1x PBS. Brains were carefully removed and then fixed in freshly prepared, ice-cold 4% paraformaldehyde (PFA, Electron Microscopy Sciences, #19210)/ PBS (pH7.4) for 4 hrs, followed by cryoprotection in the dark in 30% sucrose/0.1 % sodium azide/PBS at 4°C for at least 48 hrs. The tissue was then embedded in Tissue-Tek O.C.T Compound (Sakura Finetek, #4583) and sectioned at −22°C using a Thermo Scientific Microm HM 550. Sixteen-μm coronal brain sections were collected on Fisherbrand^™^ Superfrost^™^ Plus Microscope slides (Fisher Scientific, #12-550-15). Brain sections were further dried on the slide for 3 hrs at RT and −20°C overnight. To perform *in situ* hybridization, brain sections were first washed in 1x PBS for 5 min and then post-fixed on slides in 4% fresh PFA for 30 min. Slides were then briefly washed in RT distilled water (dH_2_O), RT 1x PBS, and then boiled in RNAscope® Target Retrieval reagent (ACDBio, #322000) at 99°C for 15 min. After a brief wash in dH_2_O, slides were incubated with freshly prepared 0.3% hydrogen peroxide for 10 min at RT, briefly washed with dH_2_O, and then incubated in 100% alcohol for 3 min. After drying at RT, brain sections were treated with RNAscope® Protease III at 40°C for 30 min, then washed with dH_2_O. Probes recognizing Adra1a mRNA (ACDBio, #408611, C1) and Pdgfra mRNA (ACDBio, #480661, C2) were diluted according to the manufacturer’s instructions and added to brain sections for 2 hrs at 40°C. Slides were then washed in 1x RNAscope® Wash Buffer (ACDBio, #310091) for 2 × 2 min. The signal of the hybridized RNA probes was amplified with RNAscope® FL v2 (ACDBio, #323110) Amp 1 for 30 min at 40°C, Amp 2 for 30 min at 40°C, and Amp 3 for 15 min at 40°C, with 2x wash in 1x Wash buffer for 2 min between each amplification step. Fluorescence signals for each color channel were then developed by first incubating slides in RNAscope® Multiplex FL v2 HRP-C1/C2 for 15 min at 40°C, then Opal Fluorophore Reagent 520/570 (Akoya Biosciences, #OP-001001/001003) mixed with 1x TSA^™^ (Tyramine Signal Amplification technology, Perkin Elmer) for 30 min at 40°C, and finally in RNAscope® Multiplex FL v2 HRP blocker (ACDBio, #323107) for 15 min 40°C. There was 2x wash in 1x Wash buffer for 2 min between each detection step. After detection steps were finished for all channels, cell nuclei were stained by DAPI for 1 min at RT and tissues were mounted in Aqua-Poly/Mount (Polysciences, #18606). Images were acquired using Zeiss LSM 880 confocal microscope with a Zeiss 20x/0.8 Plan-Apochromat objective and later analyzed using ImageJ.

### Head plate installation and cranial window surgery

The day before cranial window surgery, dexamethasone (VetOne, NDC#13985-037-02) was given through drinking water to the mice (1 mg/kg) that completed TAM administration (see above). The next day, animals were anesthetized with inhaled isoflurane (0.25-5%) and placed in a custom-made stereotaxic frame. Surgery was performed under standard and sterile conditions. After hair removal and lidocaine application (1%, VetOne, NDC 13985-222-04), the mouse’s skull surrounding the right visual cortex was exposed and the connective tissue was carefully removed. Vetbond^™^ (3M) was used to close the incision site. A custom-made metal head plate was fixed to the cleaned skull using dental cement (C&B Metabond, Parkell Inc.). A 2 × 2 mm^2^ square craniotomy was then performed using a high-speed dental drill bit. The center of the craniotomy was located 2 mm lateral to lambda. Dura was left intact and the cranial window was then sealed with a custom-made 2 × 2 mm^2^ square #1 (0.17 mm) coverslip using Vetbond^™^. A layer of cyanoacrylate (Krazy Glue) was applied on top of the Vetbond to secure the coverslip. For the LED stimulation experiment, a 014 and a 015 Viton O-ring 75A were stacked on top of the head plate with the cranial window at the center of the O-rings and glued to the head plate with black dental cement (Ortho-Jet^™^, Lang Dental). Animals recovered in their home cages for at least 4 weeks before imaging.

### In vivo 2P laser scanning microscopy for OPC Ca^2+^ activity

Two-photon laser scanning microscopy was performed with a Zeiss LSM 710 microscope equipped with a GaAsP detector, which uses a mode-locked Ti-Sapphire laser (Coherent Chameleon Ultra II) tuned to 920 nm. The power at the sample level during imaging was 60 – 100 mW depending on the depth of imaging. Higher intensities caused photoactivation of OPC Ca^2+^ activity. The head of the mouse was immobilized by attaching the head plate to a custom-made stage mounted on a vibration isolation table, and the body of the mouse was housed in a custom-made plastic restrainer. The thickness of dura was measured by autofluorescence and needed to be < 30 μm to clearly visualize OPC Ca^2+^ activity in the fine processes. Fluorescence images were collected 50 – 150 μm below dura using a coverslip-corrected Zeiss 20x/1.0 W Plan-Apochromat objective with a pixel dwelling time of 1.58 μs and scanning speed of 2 Hz. We did not detect any Ca^2+^ events shorter than 0.5 s when scanning individual processes at 10 Hz (Extended Data Fig. 7), indicating that sampling at 2 Hz adequately represented Ca^2+^ event time courses. Vascular structure and PDGFRα^+^ perivascular fibroblasts expressing GCaMP6s were used as landmarks to identify and follow individual OPCs over longitudinal imaging sessions. Mice were kept on the stage for no more than one hour and all *in vivo* imaging experiments were performed during the day. For longitudinal imaging experiments, the same cell was imaged at approximately the same time of the day. For enforced locomotion experiments, a custom-made acrylic plate was connected to an electric motor controlled by custom-made Arduino scripts and placed under a Bergamo II multiphoton microscope (Thorlabs, Inc.) equipped with a mode-locked Ti-Sapphire laser (Coherent Chameleon Discovery NX) tuned to 920 nm. Mice were head fixed during imaging under a Nikon LWD 16x/0.8 W objective with a pixel dwelling time of 1.8 μs and a frame rate of ~1.4 frames per second.

### Longitudinal imaging with 2P microscopy and spectral confocal reflectance (SCoRe) microscopy

Longitudinal imaging of OPC Ca^2+^ activity using 2P microscopy and local myelin pattern changes using SCoRe microscopy was both achieved using a Zeiss LSM 710 microscope. Fluorescent images displaying OPC Ca^2+^ activity were acquired by 2P excitation as described above, except a frequency of 1 Hz was used. To acquire myelin patterns in the cortical layer I, SCoRe microscopy was used as follows: Lasers emitting visible light of 488, 543 and 633 nm were simultaneously applied to the cortex, and light reflected by myelin at each wavelength was collected in sections of 487-490, 543-545 and 632-635 nm, respectively. Laser intensities were kept minimal to prevent tissue damage. During each imaging session, the bottom of the dura (z = 0) was determined using 2P as described above, and then a 3 min 1024 × 1024 pixel time-lapse video of OPC Ca^2+^ activity was recorded at the desired depth. Before SCoRe images were collected, the bottom of dura was re-registered as z = 0 with visible lasers to align with 2P images later during analysis. Then a 1024 × 1024 pixel z stack from the bottom of dura to at least 70 μm below the dura was collected with a step size of 2 μm. Since anesthesia dramatically disrupts OPC Ca^2+^ activity for 24 hrs (see Fig. 2), mice were awake when SCoRe microscopy was performed. Myelin structure was obtained by adding all the reflective signals collected from 487-490, 543-545 and 632-635 nm ranges. Background noise was subtracted using ImageJ (NIH). SCoRe image stacks across different days were first registered with ImageJ function Correct 3D drift and then Fijiyama was used to align myelin structure between two different time points.

### Calcium image processing and analysis

Images containing time-lapse sequences of OPC Ca^2+^ activity were first registered using moco (MOtion COrrector)^68^. The background noise was then reduced using the Kalman filter^69^. Next, images were imported into AQuA software package v1.0.1 (latest updates on Feb. 5, 2020) run under MATLAB R2018b. User-defined parameters were used to optimize the detection of OPC Ca^2+^ events and kept consistent throughout the study: Intensity threshold scaling factor: 3, Smoothing: 2, Minimum pixel size: 3, Temporal cut threshold: 3, Growing z threshold: 1, Rising time uncertainty: 2, Slowest delay in propagation: 1, Propagation smoothness: 1, Z score threshold: 2. Only processes with visible connection to the cell body were included in the analysis. Ca^2+^ events from PDGFRα^+^ perivascular fibroblasts were easily distinguishable and excluded from OPC Ca^2+^ events as the events from the former presented as either a round shape (cross section) or a tube-like structure, both without branching and propagation.

### Acute cortical slice preparation and in vitro 2P calcium imaging

Mice were deeply anesthetized with isoflurane and decapitated using a guillotine. Brains were quickly dissected out and mounted in a pre-chilled chamber on a vibratome within 5 minutes after decapitation. Cortical slices (250 μm in thickness) were sliced in ice-cold N-methyl-D-glucamine (NMDG)-based cutting solution (135 mM NMDG, 1 mM KCl, 1.2 mM KH_2_PO_4_, 1.5 mM MgCl_2_, 0.5 mM CaCl_2_, 10 mM Dextrose, and 20 mM Choline Bicarbonate, pH 7.4), and then transferred to <u>a</u>rtificial <u>c</u>erebral <u>s</u>pinal <u>f</u>luid (ACSF) (119 mM NaCl, 2.5 mM KCl, 2.5 mM CaCl_2_, 1.3 mM MgCl_2_, 1 mM NaH_2_PO_4_, 26.2 mM NaHCO_3_, 11 mM dextrose (292-298 mOsm/L)). Slices were maintained at 37°C for 40 min, and then at RT thereafter for imaging. Both the NMDG and ACSF solution were saturated with carbogen (95% O_2_ and 5% CO_2_) before use and were constantly carbogenated during the experiment. Ca^2+^ imaging was performed using Zeiss LSM 710 microscope with a Zeiss 20x/1.0 W Plan-Apochromat objective. Fluorescence images were collected at least 50 μm below the slice surface using 2P excitation at 920 nm at a frequency of 2 Hz. Registration was carried out using moco and background noise was reduced using the Kalman filter, as described above. The ΔF/F values from individual OPC were obtained by first thresholding the maximally projected movie in ImageJ, using the wand (tracing) tools to select individual OPC as a region-of-interest (ROI), and then measuring the fluorescence changes in the individual OPC/ROI.

### Visual stimulation

A LED emitting blue light was used as a light source at a distance of ~7 cm from the animal’s left eye. The light power entering the mouse left eye was ~20 nW/mm^2^. To eliminate optical cross talk between visual stimulation and 2P fluorescence detection, the objective was shielded from the light source with an opaque cylinder. The period and the interval of the visual stimulus was controlled by a pulse generator Master-8 (MicroProbes for Life Science).

### Pharmacology

For *in vivo* i.p. injection, chlorprothixene hydrochloride (Sigma-Aldrich, #C1671), prazosin hydrochloride (Tocris, #0623) and dexmedetomidine hydrochloride (Tocris, 2749) were dissolved in DMSO (Sigma-Aldrich, #D2650) at 5 mg/mL, 3 mg/mL and 0.1 mg/mL, respectively. First, 5 minutes of baseline activity was measured using 2P microscopy and then the animals were i.p. injected with equal amounts of DMSO per body weight (1 μL per 1 μg b.w.). Animals were returned to their home cage after injection and then re-mounted under the 2P microscope 20 min after injection for 5 minutes of imaging. For *in vitro* cortical slice imaging, phenylephrine hydrochloride (Tocris, #2838), tetrodotoxin citrate (Alomone labs #T-550), NBQX disodium (Tocris, #1044), (RS)-CPP (Tocris, #0173) and SR 95531 hydrobromide (Tocris, 1262) were dissolved in ACSF and applied through the superfusing solution.

### Statistical analysis

Statistical analyses were performed using OriginPro (OriginLab Corp.). All datasets had to either pass the Shapiro-Wilk test for normality before being subjected to Student’s t-test and ANOVA, or undergo non-parametric tests to determine the statistical significance. Data is reported as mean ± SEM, unless otherwise noted. P values less than 0.05 were considered statistically significant. The level of significance is marked on the figures as follows: *: p < 0.05; **: p < 0.01; ***: p < 0.001; ****: p < 0.0001; n.s.: not significant

## Supplementary Information

**Supplementary Video 1. Cytosolic OPC Ca^2+^ activity in the visual cortex *in vivo***. Left: OPC Ca^2+^ activity detected by 2P microscopy; Right: Output video from AQuA with randomly-pseudocolored Ca^2+^ events (5x speed).

**Supplementary Video 2. Membrane OPC Ca^2+^ activity in the visual cortex *in vivo***. Left: OPC Ca^2+^ activity detected by 2P microscopy; Right: Output video from AQuA with randomly-pseudocolored Ca^2+^ events (5x speed).

**Supplementary Video 3. Enforced locomotion stimulates OPC Ca^2+^ activity in the mouse visual cortex**. The red dot indicates when the platter began to rotate (5x speed).

**Supplementary Video 4. PE evokes Ca^2+^ influx in OPCs in acute cortical slices**. PE was superfused ~3 min after the recording begins (50x speed).

